# High-content imaging to phenotype antimicrobial effects on individual bacteria at scale

**DOI:** 10.1101/2021.01.11.426307

**Authors:** Sushmita Sridhar, Sally Forrest, Ben Warne, Mailis Maes, Stephen Baker, Gordon Dougan, Josefin Bartholdson Scott

**Affiliations:** Cambridge Institute of Therapeutic Immunology & Infectious Disease, University of Cambridge Department of Medicine, Jeffrey Cheah Biomedical Centre, Puddicombe Way, Cambridge Biomedical Campus, Cambridge CB2 0AW, United Kingdom; Wellcome Sanger Institute, Hinxton, CB10 1SA, United Kingdom

**Keywords:** High-content imaging, image analysis, bacteria, antimicrobial resistance, phenotyping

## Abstract

High-content imaging (HCI) is technique for screening multiple cells in high resolution to detect subtle morphological and phenotypic variation. The method has been commonly deployed on model eukaryotic cellular systems, often for screening new drugs and targets. HCI is not commonly utilised for studying bacterial populations but may be powerful tool in understanding and combatting antimicrobial resistance. Consequently, we developed a high-throughput method for phenotyping bacteria under antimicrobial exposure at the scale of individual bacterial cells. Imaging conditions were optimised on an Opera Phenix confocal microscope (Perkin-Elmer) and novel analysis pipelines were established for both Gram-negative bacilli and Gram-positive cocci. The potential of this approach was illustrated using isolates of *Klebsiella pneumoniae, Salmonella enterica* serovar Typhimurium, and *Staphylococcus aureus*. HCI enabled the detection and assessment of subtle morphological characteristics, undetectable through conventional phenotypical methods, that could reproducibly distinguish between bacteria exposed to different classes of antimicrobials with distinct modes of action (MOA). In addition, distinctive responses were observed between susceptible and resistant isolates. By phenotyping single bacterial cells, we observed intra-population differences, which may be critical in identifying persistence or emerging resistance during antimicrobial treatment. The work presented here outlines a comprehensive method for investigating morphological changes at scale in bacterial populations under specific perturbation.

**Importance:** High-content imaging (HCI) is a microscopy technique that permits the screening of multiple cells simultaneously in high resolution to detect subtle morphological and phenotypic variation. The power of this methodology is that is can generate large datasets comprised of multiple parameters taken from individual cells subjected to range of different conditions. We aimed to develop novel methods for using HCI to study bacterial cells exposed to a range of different antibiotic classes. Using an Opera Phenix confocal microscope (Perkin-Elmer) and novel analysis pipelines we created a method to study the morphological characteristics of *Klebsiella pneumoniae, Salmonella enterica* serovar Typhimurium, and *Staphylococcus aureus* when exposed to antibacterial drugs with differing modes of action. By imaging individual bacterial cells at high resolution and scale, we observed intra-population differences associated with different antibiotics. The outlined methods are highly relevant for how we begin to better understand and combat antimicrobial resistance.

## Introduction

Antimicrobial resistance (AMR) is one of the greatest current challenges in human health, with rising cases of antimicrobial resistant bacterial infections and a lack of new classes of licensed antimicrobials. Advances in bacterial genomics have revolutionised our ability to genotype antimicrobial resistant bacterial isolates at scale. However, it remains critical to link genotype with phenotype in order to interpret the biological and clinical relevance of AMR. Some phenotyping methods have been adapted to work at scale (e.g. antimicrobial susceptibility testing using semi-automated platforms such as the bioMérieux VITEK system), yet many others either rely on low throughput methods or aggregate data from mixed populations of bacterial cells. The analysis of bulk bacterial populations rather than individual cells potentially overlooks persister cells or the emergence of resistant or tolerant bacteria within that population. High throughput imaging of bacterial populations at the scale of individual cells has received limited attention but may be achieved by exploiting high-content microscopy.

High-content imaging (HCI) can be utilised as a powerful phenotypic screening approach that combines automated microscopy with image analysis to quantify multiple morphological features. This approach may capture subtle differences in structure and shape not discernible by the human eye or conventional phenotypic methods. Such image-based profiling has great potential in high-throughput drug screening, which has mainly been applied to eukaryotic cells and tissue(1, 2). In the field of microbiology, HCI has predominantly been used to study intracellular pathogens such as *Mycobacterium tuberculosis*(3–6) and *Salmonella* species as they interact with host cells(7), but only recently to screen individual bacteria growing as a population in batch culture(8, 9). Pogliano and colleagues developed a bacterial cytological profiling assay to identify morphological changes in *Escherichia coli* and other species in response to different classes of antimicrobials using fluorescence microscopy(10–13). Analysis of image data enabled the assignment of distinct morphological profiles correlating with the mechanism of action of the antimicrobial compounds tested(10). This method opened up a novel way of screening new therapeutic compounds simultaneously for efficacy and mode-of-action (MOA) using bacterial imaging(8–13).

Given the variety of AMR mechanisms harboured by bacterial species and, in many cases, by isolates of the same species, it is important to optimise HCI approaches for a range of bacteria. In this study, we developed and optimised a high-throughput imaging method based on HCI to systematically screen individual bacteria from three different species grown under antimicrobial exposure. We optimised bacterial imaging conditions using an Opera Phenix confocal microscope (Perkin Elmer) and established novel analysis pipelines for image segmentation and bacterial morphological analysis for both Gram-negative bacilli and Gram-positive cocci. The combination of HCI and image analysis enabled the detection of subtle morphological characteristics that differed between different antimicrobial classes. This work contributes to the expansion of microbial phenotyping from population-level to single-cell analysis and provides a comprehensive method of bacterial phenotypic screening at scale.

## Materials and Methods

### Bacterial isolates

A number of reference bacterial isolates, representing clinically important species, were analysed. This panel included one Gram-positive (*Staphylococcus aureus*) and two Gram-negative (*Salmonella enterica* serovar Typhimurium and *Klebsiella pneumoniae*) species. Two isolates were included per species, each with broadly different antimicrobial susceptibility profiles (Table 1).

**Table 1:** Minimum Inhibitory Concentrations.

| Species | ID | Ampicillin | Azithromycin | Trimethoprim-sulfamethoxazole | Ciprofloxacin | Gentamicin | Rifampicin | Meropenem | Tigecycline | Cefuroxime | Oxacillin | Vancomycin |
| --- | --- | --- | --- | --- | --- | --- | --- | --- | --- | --- | --- | --- |
| *K. pneumoniae* | NCTC 13438 | >256* | 32 | >32* | >32* | 6 | 16 | 0.125 | 0.094 | >256* | ND | ND |
| *K. pneumoniae* | ATCC 43816 | 64 | 2 | 0.5 | 0.16 | 1.5 | >32* | 0.125 | 0.5 | 1.5 | ND | ND |
| *S. Typhimurium* | NCTC 13347 | 0.5 | 2 | 0.25 | 0.012 | 2 | 16 | 0.125 | 0.19 | 3 | ND | ND |
| *S. Typhimurium* | NCTC 13348 | >256* | 3 | 0.25 | 0.012 | 3 | 24 | 0.064 | 0.19 | 3 | ND | ND |
| *S. aureus* | NCTC 6571 | ND | ND | 0.125 | 0.094 | 2 | ND | ND | ND | ND | 0.19 | 1 |
| *S. aureus* | ATCC 29213 | ND | ND | 0.19 | 0.25 | 2 | ND | ND | ND | ND | 0.38 | 1.5 |

MICs were determined by ETEST and are presented in µg/ml. \* indicates the MIC is above the highest antimicrobial concentration on the
ETEST; ND indicates that MIC was not determined.

### Antimicrobial susceptibility testing

Antimicrobial susceptibility testing was performed for a range of clinically relevant antimicrobials with different MOAs (Table 1). Minimum Inhibitory Concentrations (MICs) were determined by ETESTs (bioMérieux) according to the manufacturer’s instructions. Briefly, pure bacterial cultures were diluted in saline to 0.5 MacFarland standard, 100 μl of solution was inoculated and spread onto Isosensitest plates (Oxoid, CM0471), and an ETEST strip was placed on top. Plates were incubated for 16-18 hours at 37°C before the result was read.

### Preparation of plate coatings

Coating matrices were prepared according to manufacturer recommendations in sterile conditions (Table S1). All coatings, except poly-L-lysine, were incubated in ultra-thin 96 well plates (Perkin Elmer CellCarrier Ultra, 6655308) overnight at 37°C. The following day, wells were rinsed 1-3 times with wash buffer (Table S1). For poly-L-lysine, wells were coated for 5 minutes. The solution was aspirated, and wells were left to dry overnight at 37°C.

### Bacterial imaging assay

Overnight bacterial cultures were diluted in LB broth and mixed with antimicrobials to a final antimicrobial concentration of 5x MIC. Where an MIC could not be measured (i.e., where bacterial growth continued along the whole length of the ETEST), the upper limit of the ETEST was arbitrarily used in place of the MIC. The bacteria were incubated with and without antimicrobials in static incubators in ultra-thin 96 well plates for 2 hours at 37°C. The plates were aspirated, and the remaining adherent bacteria were fixed with 4% paraformaldehyde (Alfa Aesar, J61899.AK) for 10 minutes. The wells were washed once with 50μl of DPBS (Thermo, 10010023) before staining. Fixed cells were stained with FM4-64 (2μg/ml, Thermo, T13320), SYTOX green (0.25μM, Thermo, S7020) and 4′,6-Diamidino-2-phenylindole dihydrochloride (DAPI, 2μg/ml, Sigma, D9542). Staining was performed at ambient temperature for 20 minutes in the dark followed by a wash with 50μl PBS. Finally, 50μl of PBS was added to wells and the plates were imaged within 24 hours.

### High-content imaging and image analysis

High-content confocal imaging was performed using an Opera Phenix (Perkin Elmer), using a 63x water immersion lens. 10 fields of view (equating to 0.4 mm^2^) were imaged for each well, with 3 z-stacks per field at 0.5 μm intervals to ensure comprehensive imaging of the bacterial monolayer. Triplicate biological and technical replicates were performed for all experiments. Image analysis was performed using Harmony (v4.9). Optical correction was performed using flatfield and brightfield correction. The detailed full analysis pipelines are shown in Table S1 and S2. Data were exported and plotted in GraphPad Prism and R(14).

## Results

### High-content imaging and analysis of individual bacteria

An HCI workflow was established using two reference isolates from each of the three bacterial pathogens *S*. Typhimurium, *K. pneumoniae*, and *S. aureus* (Figure 1A; Table 1). Organisms were selected to have contrasting AMR profiles within each species. Each of the isolates was exposed to different antimicrobial agents, and HCI was used to collect phenotypic data for numerous individual bacteria within each assay. To this end, overnight bacterial cultures were grown in 96-well microtiter plates in the presence or absence of each antimicrobial for two hours to capture multiple early morphological changes. The antimicrobials used are listed in Table 1. To capture images, the bacteria in each well were stained *in situ* with markers for the cell membrane (FM4-64), nucleic acid (DAPI), and membrane permeability (SYTOX green)(10) (Figure 1B). Imaging was performed on an Opera Phenix and image analysis was conducted using Harmony software.

**Figure 1:**
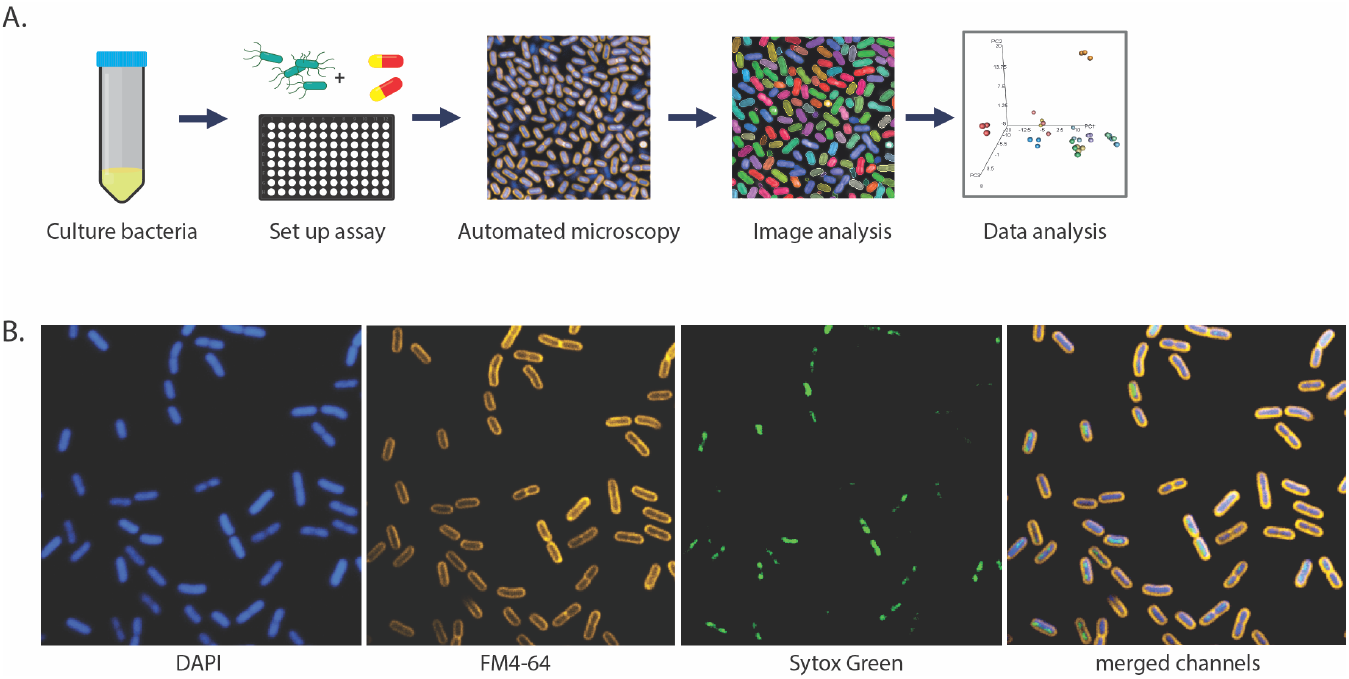
Bacterial high-content imaging. (**A**) Schematic of the bacterial high-content imaging workflow. Overnight bacterial cultures are added to ultra-thin bottom plates, and incubated, with or without antimicrobial compounds. Adherent bacteria are fixed and stained before being imaged on an Opera Phenix high-content confocal microscope using a 63x water immersion objective. Images were analysed using Harmony software and data was exported and plotted in R. (**B**) Representative image of *K. pneumoniae* NCTC 43816 stained with FM4-64 (cell membrane), DAPI (nucleic acid, membrane permeable) and SYTOX green (nucleic acid, membrane impermeable).

As both rod- and cocci-shaped bacteria were imaged at the single cell scale, it was necessary to build separate, parallel pipelines for accurate analysis of microorganisms with different morphology. Examples of image segmentation and analysis of Gram-negative rods and Gram-positive cocci are shown in Figure 2 and detailed in Table S2 and Table S3, respectively. Images of Gram-negative rods were initially filtered using FM4-64 intensity patterns to enhance stained objects and subtract the background (Figure 2A). All images were then calculated based on DAPI and FM4-64 intensity and resized to include both cytosolic (DNA) and membrane regions; filtering was performed to remove artefacts such as incomplete bacterial bodies (Figure 2A and C). Morphological features and stain intensities were calculated for each defined bacterial cell, including area, roundness, width and length, as well as various measures of the intensity, symmetry and distribution of each of the FM4-64, DAPI and SYTOX green channels within each object (Figure 2B and D). For Gram-negative rods, the segmented objects were further classified as either single bacterial cells, dividing cells, or artefacts (Figure 2B) with a manually trained linear classifier using Harmony PhenoLOGIC. Subsequent analyses were conducted on the single cells only. The number of individual bacteria captured and analysed per well was dependent on isolate and treatment but was a minimum of 2,000 bacteria per untreated well for all replicates.

**Figure 2:**
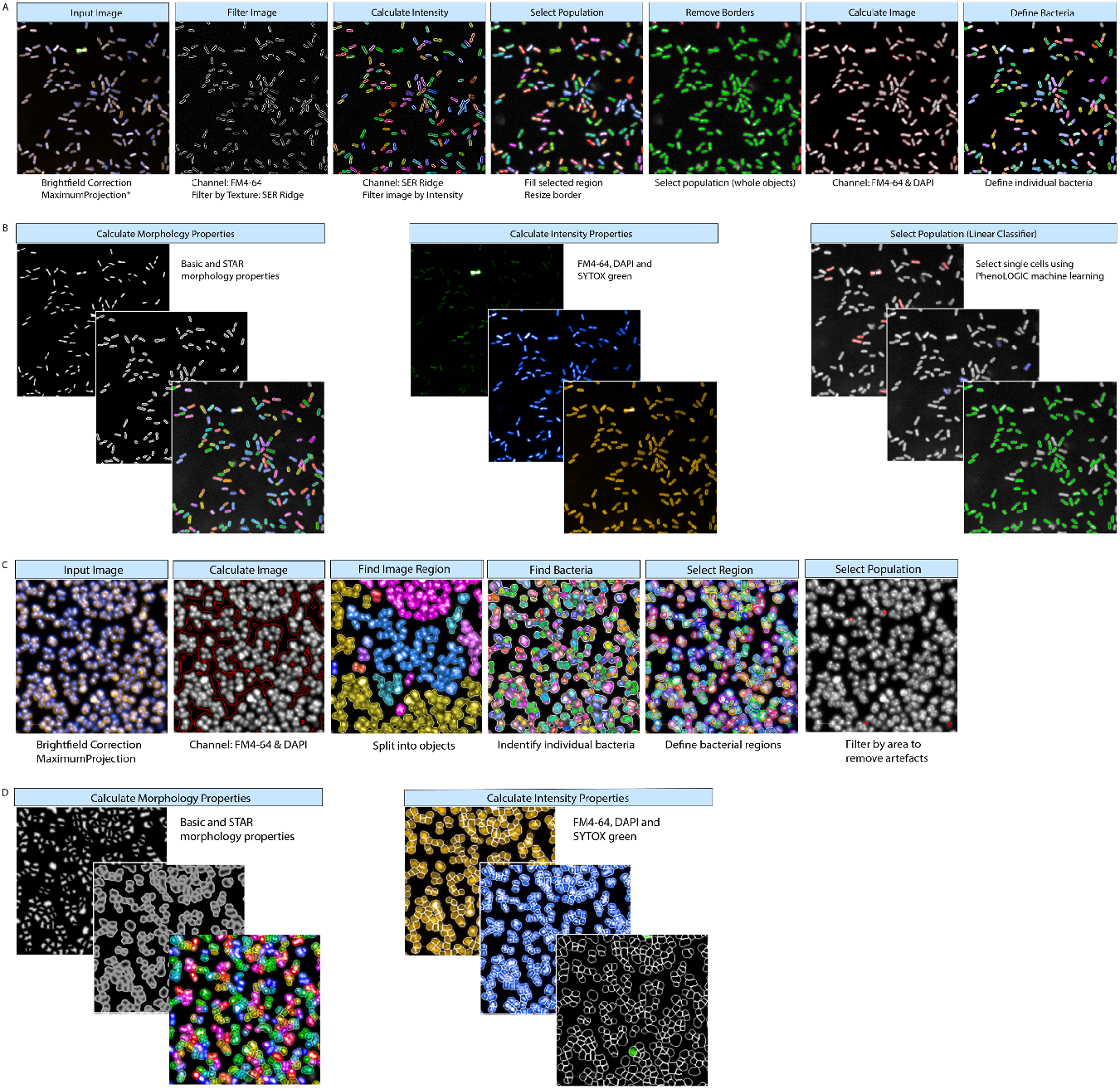
Harmony bacterial image analysis workflow for Gram-negative rods (A-B) and Gram-positive cocci (C-D). (**A**) Using basic brightfield correction and maximum projection, images were segmented by filtering the images using texture properties based on the FM4-64 channel to remove any background. The image region was filled and resized and border objects were excluded to include only whole objects. The image region was further calculated using FM4-64 and DAPI fluorescence and individual bacteria were defined. (*single planes were analysed for *S.* Typhimurium) (**B**) Bacterial morphology and stain intensity properties were calculated using DAPI, SYTOX Green and FM4-64 fluorescence. Finally, a linear classifier was used to train the software to define single bacterial cells and exclude any artefacts. (**C**) Using basic brightfield correction and maximum projection, the bacterial region was defined using a calculated image based on DAPI and FM4-64 channels. Individual bacteria were identified within the image region, and the bacterial regions were defined and resized into individual bacterial cells. Any artefacts were removed using size filters. (**D**) Bacterial morphology and stain intensity properties were calculated using DAPI, SYTOX Green and FM4-64 fluorescence.

### Optimising bacterial imaging

The bacterial isolates displayed different degrees of adhesion to the base of the 96-well plates, which significantly affected the image quality and downstream analysis. For example, the non-motile *K. pneumoniae* isolates spontaneously strongly adhered to the bottom of the wells, whereas the motile *S.* Typhimurium isolates displayed relatively poor adhesion resulting in blurry superimposed images of the flattened z-stack (maximum projection) (Figure S1A and B). Consequently, it was necessary to assess multiple plate coating conditions for each *S.* Typhimurium isolate to identify the optimal conditions for binding and image clarity. Ultimately, the image segmentation pipelines were used to quantify individual bacteria on 11 commercially available coating matrices (thick and thin rat tail collagen, Matrigel, vitronectin, fibronectin, Cell-Tak, laminin, wheat germ agglutinin (WGA), poly-L-lysine, gelatine, and mouse collagen) in comparison to non-coated wells (Table S1).

The optimal coating conditions were found to differ between species, and to a lesser extent for each isolate (Figure 3). The *K. pneumoniae* isolates displayed the best adhesion and image quality on non-coated, WGA, and fibronectin-coated plates (Figure 3A, Figure S2A). Therefore, all subsequent experiments with these isolates were conducted on non-coated wells. In contrast, image quality and the number of adherent *S.* Typhimurium bacteria improved dramatically upon optimization of the plate coating (Figure S1C). Both *S.* Typhimurium isolates displayed very poor adhesion to non-coated wells (Figure 3B; Figure S2B), but adhered sufficiently, although to different extents, to wells coated with thick rat tail collagen, Matrigel, and vitronectin (Figure 3B; Figure S2B), with collagen and vitronectin being the optimal conditions for NCTC 13347 and NCTC 13348 adhesion, respectively. While different coatings were chosen for the two isolates, it would have been feasible to use the same coating, as the number and image quality of adhered organisms on rat tail collagen, Matrigel, and vitronectin were sufficient for analysis. To overcome any residual lack of adhesion of *S.* Typhimurium, image analysis of these isolates was performed on individual planes rather than a maximum projection of three z-stacks.

**Figure 3:**
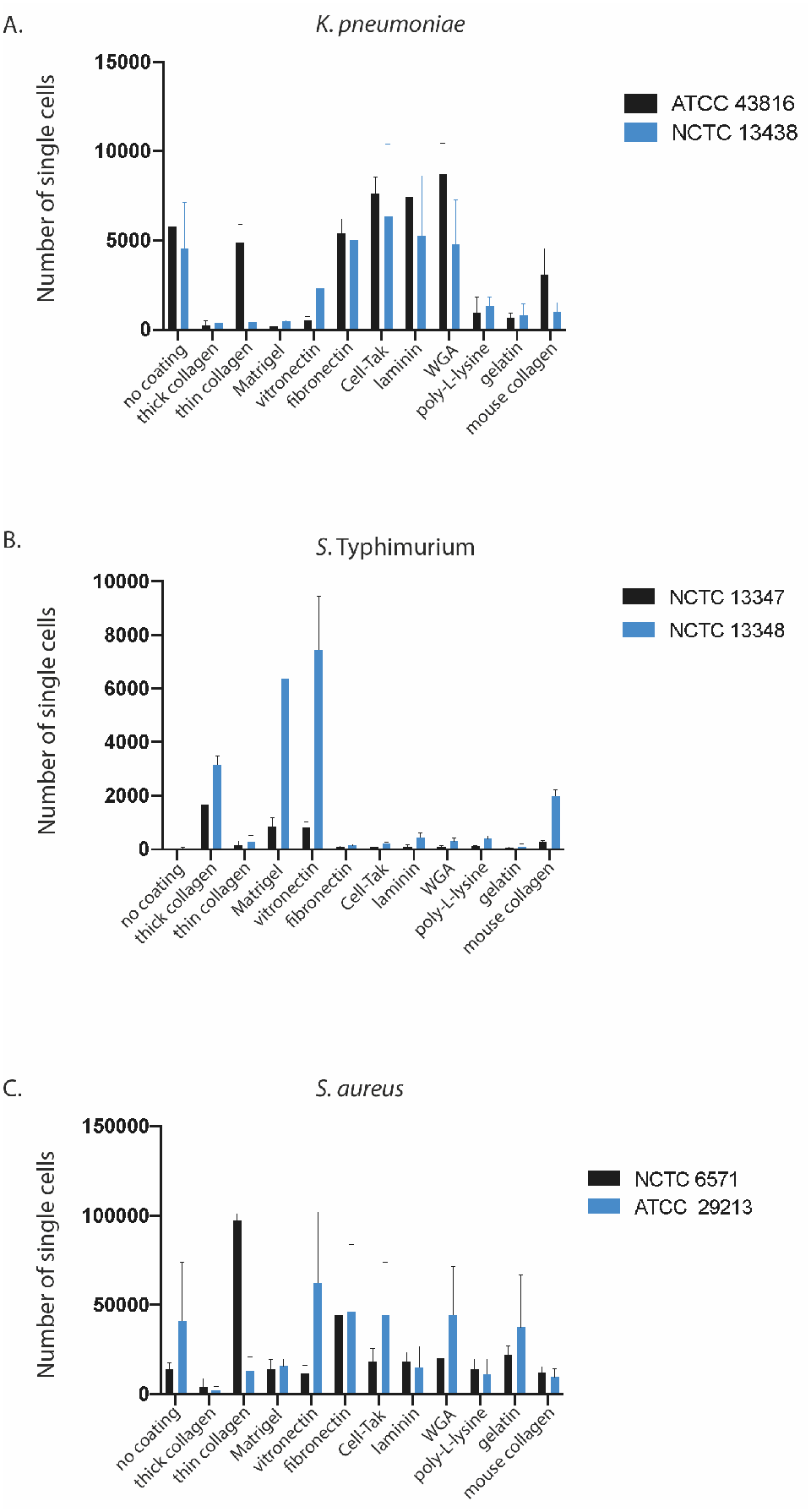
Optimising plate coating for bacterial adhesion. Isolates were grown in ultra-thin 96 well plates on different surface matrices and the Harmony analysis pipelines were used to count the number of adherent bacteria after fixing, washing and staining. Graphs are comparing the adhesion of two representative isolates of *K. pneumoniae* (A), *S.* Typhimurium (B) and *S. aureus* (C) on each substrate. Error bars represent standard deviation of three biological replicates.

The two *S. aureus* isolates displayed different adhesion properties, with ATCC 29213 showing increased adhesion on vitronectin, fibronectin, Cell-Tak, WGA, and gelatin-coated wells, whereas NCTC 6571 only had sufficient cell counts on thin collagen and fibronectin coated wells (Figure 3C). Taking adhesion and image clarity (Figure S2C) into account, thin collagen and vitronectin were used for optimal adhesion of NCTC 6571 and NCTC 29213, respectively.

### Measuring distinct morphological changes in response to antimicrobial compounds

To measure the phenotypic effects of antimicrobials with distinct mechanisms of action (MOAs), bacteria were incubated with 11 commercially available antimicrobials for 2 hours and imaged as described above. Antimicrobials were used at 5x the MIC determined by ETEST, or 5x the highest concentration tested if an isolate had an MIC higher than the ETEST range (Table 1). Figure 4A, 5A, and 6A show examples of the observed morphological changes at 2h post-treatment for a non-motile (*K. pneumoniae* NCTC 43816) and a motile Gram-negative rod (*S.* Typhimurium NCTC 13348), and a Gram-positive coccus (*S. aureus* ATCC 29213), respectively.

**Figure 4:**
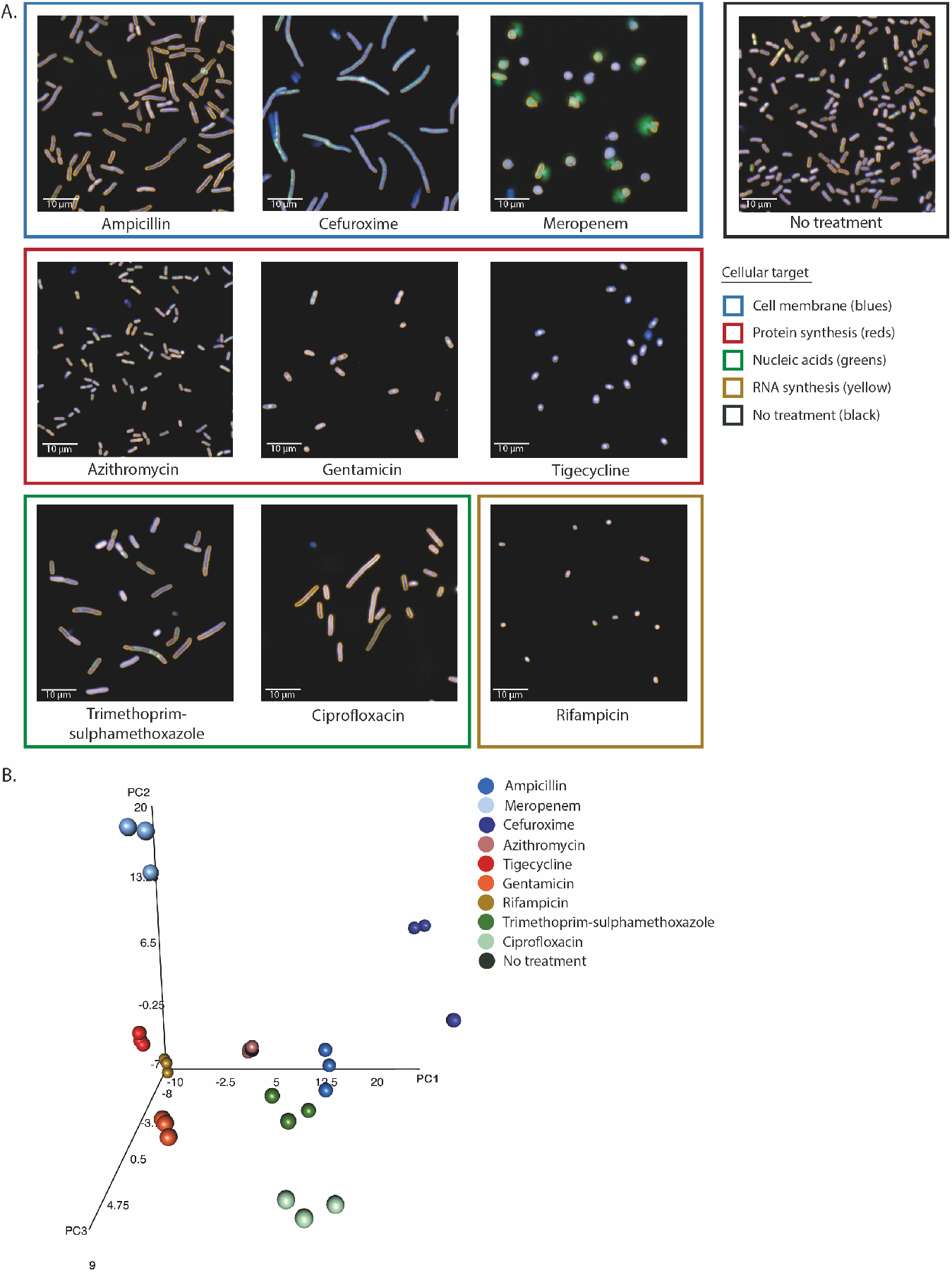
Morphological effects on *K. pneumoniae* NCTC 43816 under antimicrobial pressure. (**A**) Representative images of the effect of different antimicrobials on the *K. pneumoniae* isolate NCTC 43816 in exponential growth phase after 2 hours of incubation. Antimicrobials are grouped by similar cellular targets. Bacteria were stained with FM4-64, DAPI and SYTOX green. Images were acquired on an Opera Phenix using a 63x water immersion lens. (**B**) Three-dimensional principal component analysis of the mean and standard deviation values of 62 morphological properties measured for single bacterial cells in each well. Technical triplicate repeats are shown.

**Figure 5:**
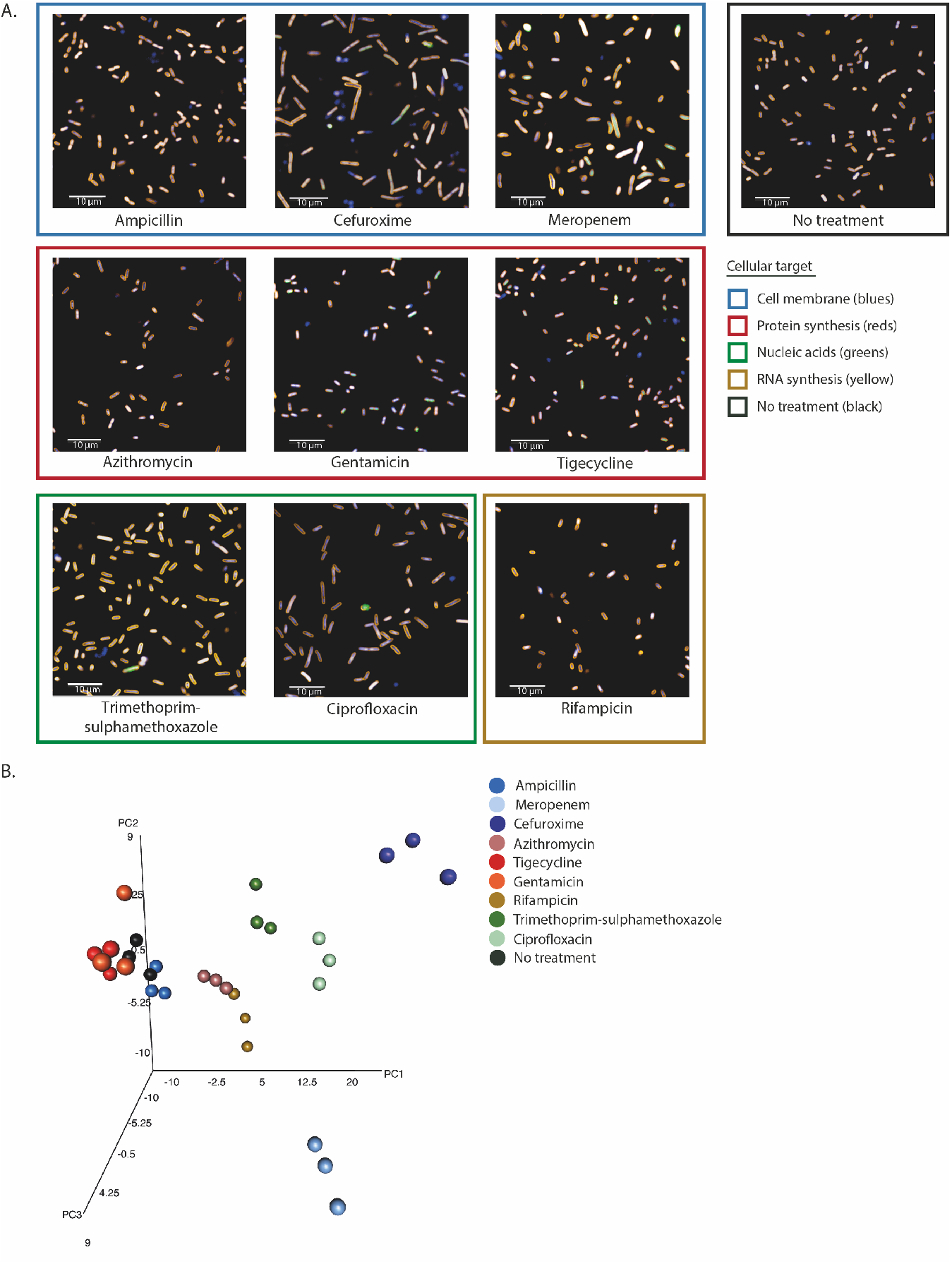
Morphological effects on *S.* Typhimurium NCTC 13348 under antimicrobial pressure. (**A**) Representative images of the effect of different antimicrobials on the *S.* Typhimurium isolate NCTC 13348 in exponential growth phase after 2 hours of incubation. Antimicrobials are grouped by similar cellular targets. Bacteria were stained with FM4-64, DAPI and SYTOX green. Images were acquired on an Opera Phenix using a 63x water immersion lens. (**B**) Three-dimensional principal component analysis of the mean and standard deviation values of 62 morphological properties measured for single bacterial cells in each well. Technical triplicate repeats are shown.

**Figure 6:**
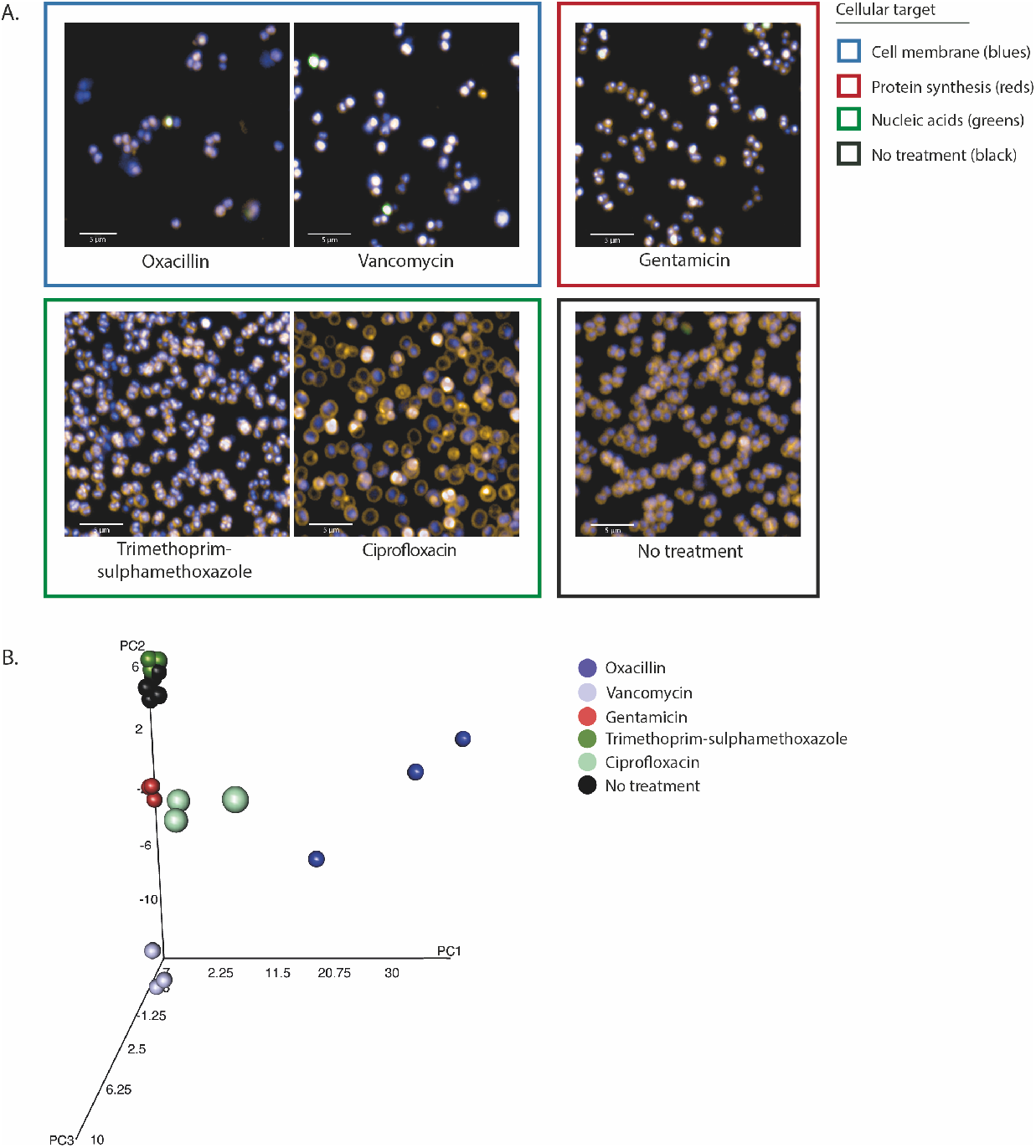
Morphological effects on *S. aureus* ATCC 29213 under antimicrobial pressure. (**A**) Representative images of the effect of different antimicrobials on the *S. aureus* isolate ATCC 29213 in exponential growth phase after 2 hours of incubation. Antimicrobials are grouped by similar cellular targets. Bacteria were stained with FM4-64, DAPI and SYTOX green. Images were acquired on an Opera Phenix using a 63x water immersion lens. (**B**) Three-dimensional principal component analysis of the mean and standard deviation values of 62 morphological properties measured for single bacterial cells in each well. Technical triplicate repeats are shown.

There were notable differences that the pipeline was able to capture between the effects of the same antimicrobials on Gram-negative versus Gram-positive bacteria, with more visually striking morphological changes observed in the Gram-negative bacteria. The established image analysis pipelines produced mean and standard deviation measurements for >90 morphological features and stain intensities for each bacterium imaged (Table S4-6). These measurements were combined for each isolate and analysed using principal component analysis (PCA). Technical replicates of each class of antimicrobial separated into distinct clusters based on MOA (Figure 4B, 5B and 6B, Figure S3), and biological replicates produced similar distribution by PCA, demonstrating assay reproducibility (Figure S4). Although separate pipelines were required for Gram-positive and Gram-negative organisms, each pipeline was able to distinguish a wide variety of phenotypes generated by antimicrobial treatment, segmenting the images and identifying individual bacteria despite the morphological changes associated with each antimicrobial (Figure S5).

Antimicrobials acting on similar cellular processes generally induced comparable morphological changes in each species, and these were found to cluster in a PCA. Bacteria treated with tigecycline and gentamicin, which block protein synthesis by binding the 30S ribosomal subunit, clustered proximally for all Gram-negative isolates tested (Figure 4B, 5B). In addition, these generally also clustered near rifampicin and azithromycin treated bacteria (Fig 4B, 5B); these antimicrobials affect protein synthesis by inhibiting RNA polymerase or translation by binding the 50S ribosomal subunit, respectively. Antimicrobials that inhibit DNA synthesis (trimethoprim/sulphamethoxazole), DNA replication (ciprofloxacin) and cell wall synthesis (ampicillin, cefuroxime and meropenem) tended to induce an elongated phenotype and again clustered proximally (Fig 4B, 5B). Notably, meropenem clustered separately to the other b-lactams for the *Klebsiella* isolates and appeared to disrupt the bacterial cell wall more potently, causing the bacteria to swell and lyse instead of elongating (Figure 4, S3A). *K. pneumonia* NCTC 13438 was resistant to ampicillin, trimethoprim/sulphamethoxazole, cefuroxime and ciprofloxacin at concentrations higher than the ETEST scale, and with the exception of ciprofloxacin, these clustered with the untreated control (Figure S3A). A similar phenotype was observed for *S.* Typhimurium NCTC 13348 treated with ampicillin (Figure 5B). This highlights that HCI screens provide novel data regarding drug susceptibility as well as MOA.

The morphological changes observed for *S. aureus* were relatively subtle compared to those for the Gram-negative isolates (Figure 6A). Only ciprofloxacin induced a visually discernible phenotypic change, which was associated with enlarging bacterial area. However, after image analysis, each antimicrobial effectively separated into unique clusters by PCA, except trimethoprim/sulphamethoxazole, which clustered alongside the untreated controls. (Figure 6B, S3C, S5). This finding suggests that the analysis could discriminate between very subtle cellular variation by capturing and analysing a large number of phenotypic parameters.

### Measuring the relative importance of specific morphological and fluorescence intensity parameters

To assess the quality of the image analysis we calculated the Z prime (Z’) values using Harmony, comparing treated and untreated bacteria for each species and antimicrobial combinations (Table S7-9), where an ideal assay should yield values between 0.5 and 1(15). The Z’ values were higher for antimicrobials that clustered further from the untreated control in the PCA, and the Gram-negative isolates generally had more Z’ values above 0.5 than the Gram-positive isolates. For example, the trimethoprim/sulphamethoxazole-treated *S. aureus* ATCC 29213 failed to separate from the untreated control by PCA, which correlated with poor Z’ values (Table S9). Similarly, poor Z’ values were obtained for azithromycin-treated *K. pneumoniae* NCTC 43816 (which is intrinsically resistant to macrolides) and gentamicin-treated *S*. Typhimurium NCTC 13348 (Table S7-8).

To highlight the relative importance of some of the measured parameters with high Z’ values, differences in measurements across representative antimicrobials were compared.

Morphological measurements of roundness, area and length-to-width ratio (Figure 7), as well as threshold compactness and the radial relative deviation of the DAPI and FM4-64 staining patterns, were plotted for a selection of antimicrobials with different MOAs (Figure S6). In addition, SYTOX green intensity was included as this should only stain bacteria if membrane integrity has been compromised (Figure S6 M-O).

**Figure 7:**
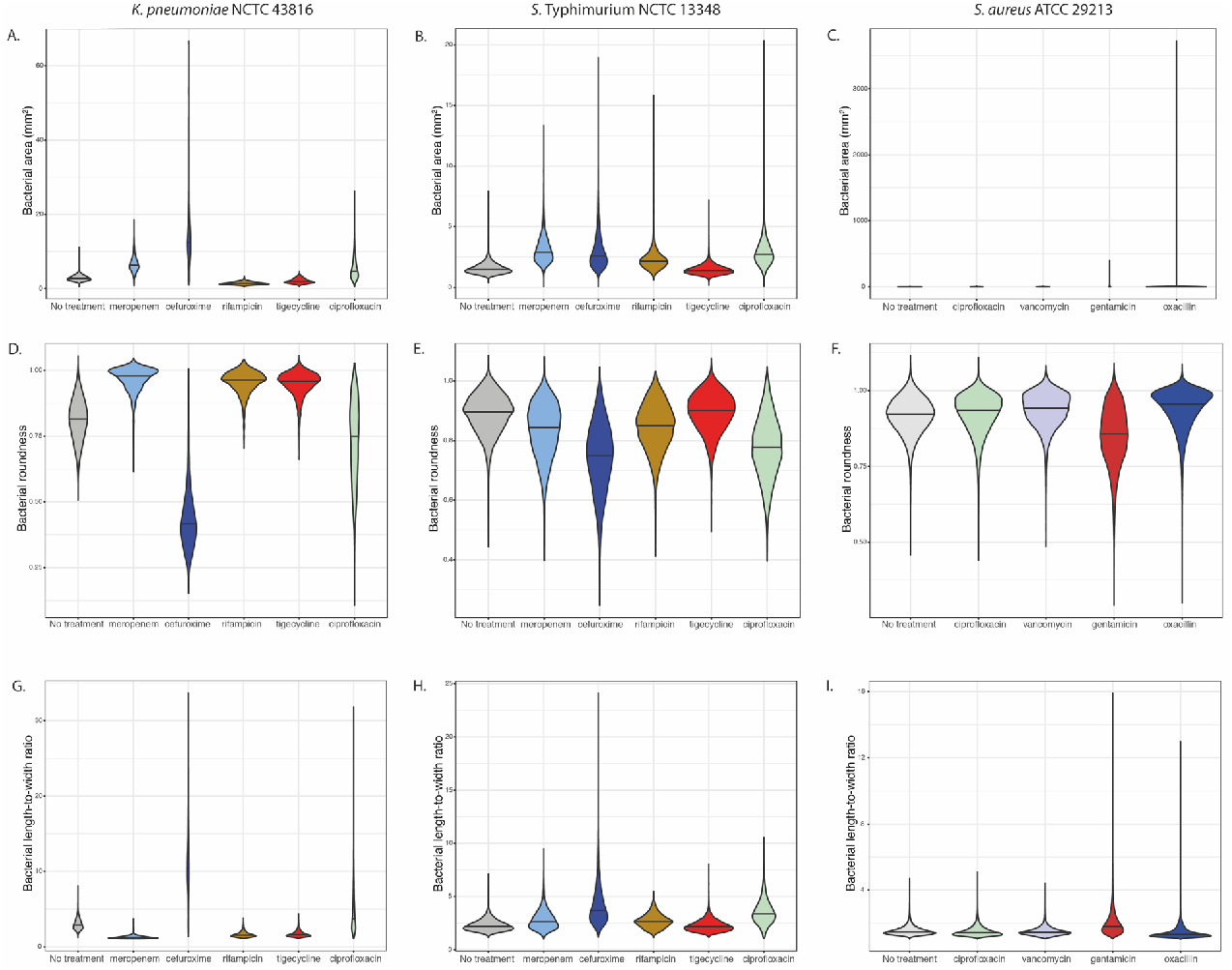
Comparison of individual basic morphological measurements. Violin plots of bacterial area (A-C), bacterial roundness (D-F) and bacterial length-to-width ratio (G-I) comparing *K. pneumoniae* NCTC 43816 and *S.* Typhimurium NCTC 13348 treated with meropenem, cefuroxime, rifampicin, tigecycline and ciprofloxacin, and *S. aureus* ATCC29213 treated with ciprofloxacin, vancomycin, gentamicin and oxacillin, with untreated controls.

When plotting these parameters individually, clear differences were observed between the different antimicrobials for the *K. pneumoniae* isolates and, to a lesser extent, the *S.* Typhimurium isolates. For example, increased area, decreased roundness, increased length-to-width ratio and FM4-64 and DAPI radial relative deviation correlated with the observed elongation phenotype observed for cefuroxime and ciprofloxacin (Figure 7 A, B, D, E, G, H and Figure S6 A, B, D, E). In contrast, increased FM4-64 and DAPI threshold compactness as well as bacterial roundness was observed for rifampicin and tigecycline (Figure 7 D, E, and Figure S6 G, H, J, K).

Generally, SYTOX green intensity was higher for antimicrobials disrupting the bacterial membrane for both Gram-negative (meropenem) and Gram-positive isolates (oxacillin). However, the effect of individual parameters on *S. aureus* were subtler than for the Gram-negative isolates, with only gentamicin treatment showing slightly decreased roundness and increased length-to width ratio (Figure 7 F and I), demonstrating the need to observe multiple combined phenotypic parameters.

### Phenotypes within a bacterial population

Using violin plots, it was possible to visualize the population density and distribution of bacteria for any given parameter. This analysis demonstrated the inherent morphological heterogeneity in bacterial populations of the same isolate under the same growth conditions (Figure 7 and S6). For example, certain antimicrobial treatments yielded a high degree of heterogeneity in the length-to-width ratio, notably cefuroxime and ciprofloxacin treated *K. pneumoniae* NTCT 43816 (Figure 8). In contrast, rifampicin treatment appeared to yield decreased variability within a population as compared to untreated controls (Figure 8). By phenotyping single cells, it is possible to observe within-population differences, which is critical for identifying persistence or emerging resistance during antimicrobial treatment.

**Figure 8:**
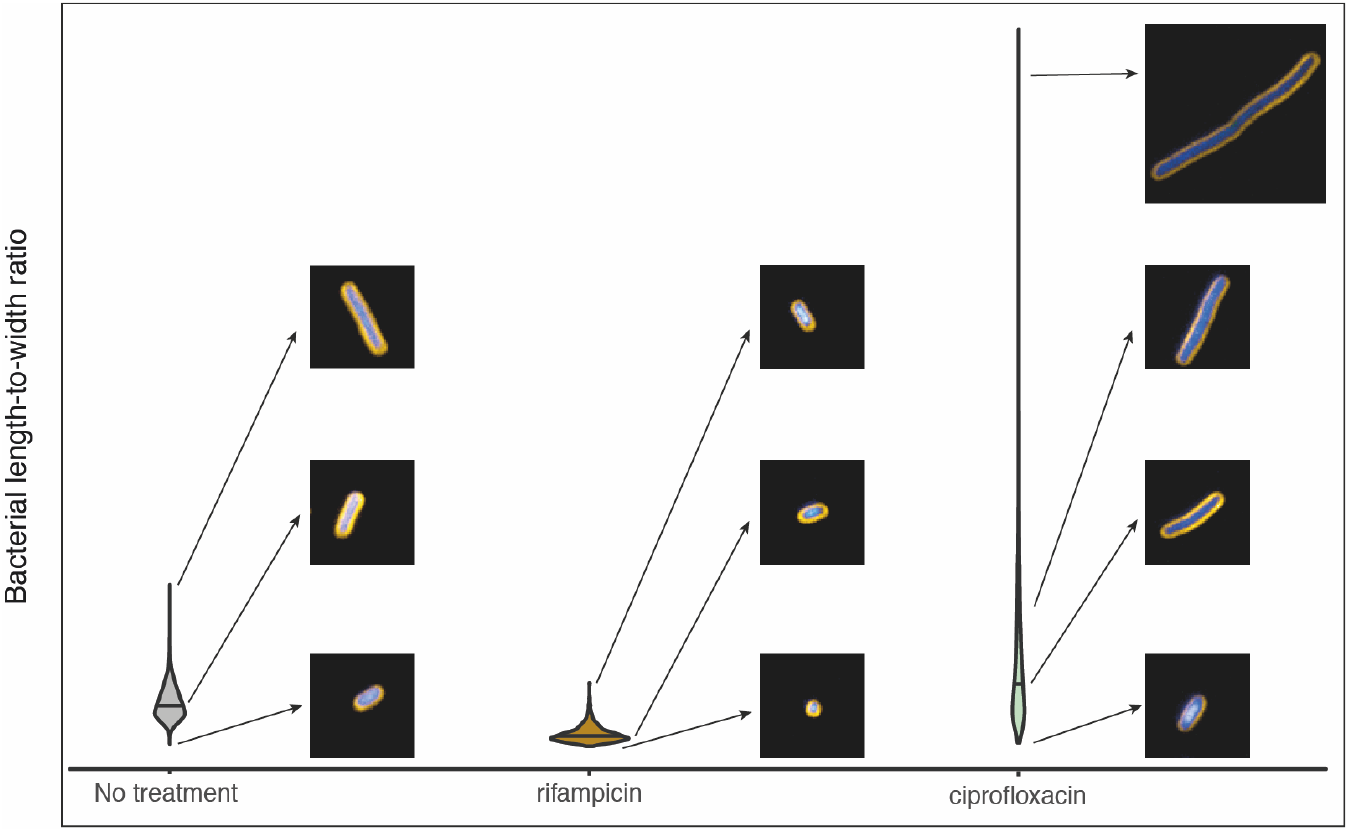
Example of population-level length heterogeneity of *K. pneumoniae* NCTC 43816. Violin plot of bacterial length-to-width ratio comparing untreated, rifampicin treated, and ciprofloxacin treated *K. pneumoniae* NCTC 43816 with inset images demonstrating the different phenotypes observed in the same growth conditions within a single well.

## Discussion

In this study we optimised an experimental pipeline for high-throughput confocal imaging of motile and non-motile bacteria in liquid culture. We used this method for systematic screening of Gram-positive and Gram-negative bacteria under antimicrobial pressure, with robust and standardised image analysis pipelines to efficiently and reproducibly measure distinct morphological changes correlating with antimicrobial MOA. This analysis was built around profiling the subtle morphological phenotypes of individual bacteria in a culture, providing information on the whole population and variation within that population.

There is a number of advantages to using HCI for bacterial research. It provides flexibility in experimental design, with the ability to customise and compare growth conditions and individual isolates from different species in high throughput. Traditional phenotyping methods rely on the collective properties of large numbers of bacteria, HCI enables measurements at the scale of individual bacterial cells. Advances in image analysis permit reliable segmentation of bacterial images and rapid, detailed profiling of individual bacterial cells with the ability to demonstrate the heterogeneity of bacterial phenotypes in any given environment.

Our work identified some challenges in using HCI for bacterial research, in particular variation in adhesion to microtiter plates. Poor adhesion influences both image quality and the number of bacteria successfully imaged for downstream analysis. This challenge was overcome by testing a range of coating matrices, which demonstrated substantial inter- and some intra-species variation in their ability to adhere to each substrate. For example, there were notable differences in adhesion between the non-motile *K. pneumoniae* and motile *S.* Typhimurium. *K. pneumoniae* possess an array of adhesins that allow them to adhere and persist in different environments, which have contributed to their emergence as an important nosocomial pathogen(16, 17). In contrast, *S.* Typhimurium relies on motility and more specific cellular interactions and invasion for causing infection(7). These factors highlight the need to optimise imaging conditions for each bacterial isolate. However, for most isolates, more than one coating condition was sufficient for downstream analysis, making it possible to screen multiple isolates in parallel using the same plate coating for the higher throughput assays.

One of the most challenging aspects of image analysis was the segmentation and identification of individual bacteria. This is in part because most existing image analysis software is designed primarily to analyse images of eukaryotic cells. However, analysis pipelines to effectively segment both rod- and cocci-shaped bacteria were created using existing image analysis tools in the Harmony software. Though the analysis pipelines in this study were created using Harmony, which is a proprietary software from Perkin Elmer, there are open access image analysis software options available - for example CellProfiler(18) and Cellpose(19)– which have similar analysis capabilities.

It was necessary to produce separate pipelines for cocci and rod-shaped bacteria for the initial segmentation. Other studies have also utilised different pipelines for phenotypically variant species; for example, the analysis used by Zoffmann and colleagues for *E. coli* was not suitable for *Acinetobacter baumannii*, as these species differ in size and shape(8). Importantly, the pipelines created in our study could be used to reproducibly segment bacteria in all growth conditions used, even as morphologies changed due to antimicrobial exposure. Distinct morphological changes were observed in response to different classes of antimicrobials, with different effects observed in Gram-negative versus Gram-positive species. However, bacteria from the same species generally displayed similar morphological distributions by principal component analysis when treated with 5x the MIC, correlating with antimicrobial mechanism. In addition, different clustering was observed between susceptible and resistant isolates, allowing for simultaneous evaluation of potency as well as MOA.

The phenotypic changes identified in this study in the presence of antimicrobials are comparable to previous imaging studies in *Enterobacteriaceae*, including bacterial enlargement with carbapenems and cephalosporins(20, 21), compaction of the nucleoid with antimicrobials targeting the bacterial ribosome(22) and filamentous elongation in the presence of fluoroquinolones(23). In agreement with other studies, we identified similar morphological changes in isolates of *K. pneumoniae* and *S.* Typhimurium to those previously reported for *E. coli* in response to a range of antimicrobial classes(10), but here we employed a simplified method by removing centrifugation steps and by imaging directly in wells rather than on agarose pads. This facilitates higher throughput and scalability.

The fluorescent staining protocol previously optimised by Nonejuie *et al.*(10) worked well across all the isolates tested in this study. FM4-64 stains the cell membrane, and the staining patterns should relate to membrane integrity. DAPI and SYTOX green both stain nucleic acids, but only DAPI is permeable through an intact cell membrane, making SYTOX green intensity an additional measurement of membrane integrity after antimicrobial exposure(24, 25). In addition, nucleic acid stains can distinguish between subtle alterations in nucleic acid distribution patterns. Plotting individual phenotypic parameters was sufficient when an antimicrobial induced a strong visual phenotypic effect, for example, length-to-width ratio could be used for ciprofloxacin or cefuroxime treated *K. pneumoniae*. However, in most cases, and in particular for the smaller cocci-shaped *S. aureus* isolates where the phenotypic effects were subtler, a combination of morphological as well as stain intensities, distribution and symmetry measurements were required to efficiently evaluate the data. This highlights that the software can detect important variations that are not obvious in conventional phenotypic methods.

Our methods contribute to moving microbial phenotyping from a population-based analysis to the scale of individual bacterial cells and provides a comprehensive method of bacterial phenotypic screening at scale. This approach has a wide range of applications, but the ability to provide analysis of diverse collections of isolates simultaneously in a range of growth conditions gives it important potential in the fight against AMR. In addition to existing roles in compound screening for antimicrobial efficacy and simultaneous MOA prediction(26), the technology could be used for more detailed mechanistic follow up studies using mutant libraries to assess genes that are protective against individual drugs(27). Large numbers of compounds and bacterial isolates, representing species with diverse genetic backgrounds, can be screened at scale. We have previously shown the utility of bacterial HCI for therapeutic antibody screening(9), and there is potential to assess synergy between antimicrobials and monoclonal antibodies against multi-drug resistant bacteria that would be challenging using other platforms. Importantly, by analysing individual bacteria within a culture, it is possible to detect differential effects and persister cells during drug treatment and be able to truly evaluate the efficacy of a compound.

## Acknowledgments

We thank James Hutt and Achim Kirsch for their help with the analysis pipelines. This work was supported by a Innovate UK Commercial in Confidence grant to purchase the Opera Phenix. SS and SB are funded by the Wellcome Trust (206194 and 215515/Z/19/Z). SF, BW, MM, SB, GD and SJB are supported by funding from the National Institute for Health Research [Cambridge Biomedical Research Centre at the Cambridge University Hospitals NHS Foundation Trust] and National Institute for Health Research AMR Research Capital Funding Scheme [NIHR200640]. The views expressed are those of the authors and not necessarily those of the NHS, the NIHR or the Department of Health and Social Care.

## Data Availability

All data underlying the results are available in the supplementary files associated with the article.

## References

1. van Vliet E, Daneshian M, Beilmann M, Davies A, Fava E, Fleck R, Julé Y, Kansy M, Kustermann S, Macko P, Mundy WR, Roth A, Shah I, Uteng M, van de Water B, Hartung T, Leist M. 2014. Current approaches and future role of high content imaging in safety sciences and drug discovery. ALTEX 31:479–493.

2. Bray M-A, Singh S, Han H, Davis CT, Borgeson B, Hartland C, Kost-Alimova M, Gustafsdottir SM, Gibson CC, Carpenter AE. 2016. Cell Painting, a high-content image-based assay for morphological profiling using multiplexed fluorescent dyes. Nat Protoc 11:1757–1774.

3. Christophe T, Ewann F, Jeon HK, Cechetto J, Brodin P. 2010. High-content imaging of Mycobacterium tuberculosis-infected macrophages: an in vitro model for tuberculosis drug discovery. Future Med Chem 2:1283–1293.

4. Barczak AK, Avraham R, Singh S, Luo SS, Zhang WR, Bray MA, Hinman AE, Thompson M, Nietupski RM, Golas A, Montgomery P, Fitzgerald M, Smith RS, White DW, Tischler AD, Carpenter AE, Hung DT. 2017. Systematic, multiparametric analysis of Mycobacterium tuberculosis intracellular infection offers insight into coordinated virulence. PLoS Pathog 13:1–27.

5. Manning AJ, Ovechkina Y, McGillivray A, Flint L, Roberts DM, Parish T. 2017. A high content microscopy assay to determine drug activity against intracellular Mycobacterium tuberculosis. Methods 127:3–11.

6. Greenwood DJ, Dos Santos MS, Huang S, Russell MRG, Collinson LM, MacRae JI, West A, Jiang H, Gutierrez MG. 2019. Subcellular antibiotic visualization reveals a dynamic drug reservoir in infected macrophages. Science (80-) 364:1279–1282.

7. Antoniou AN, Powis SJ, Kriston-Vizi J. 2019. High-content screening image dataset and quantitative image analysis of Salmonella infected human cells. BMC Res Notes 12:1–4.

8. Zoffmann S, Vercruysse M, Benmansour F, Maunz A, Wolf L, Blum Marti R, Heckel T, Ding H, Truong HH, Prummer M, Schmucki R, Mason CS, Bradley K, Jacob AI, Lerner C, Araujo del Rosario A, Burcin M, Amrein KE, Prunotto M. 2019. Machine learning-powered antibiotics phenotypic drug discovery. Sci Rep 9:1–14.

9. Maes M, Dyson ZA, Smith SE, Goulding DA, Ludden C, Baker S, Kellam P, Reece ST, Dougan G, Scott JB. 2020. A novel therapeutic antibody screening method using bacterial high-content imaging reveals functional antibody binding phenotypes of Escherichia coli ST131. bioRxiv 2020.05.22.110148.

10. Nonejuie P, Burkart M, Pogliano K, Pogliano J. 2013. Bacterial cytological profiling rapidly identifies the cellular pathways targeted by antibacterial molecules. Proc Natl Acad Sci 110:16169–16174.

11. Quach DT, Sakoulas G, Nizet V, Pogliano J, Pogliano K. 2016. Bacterial Cytological Profiling (BCP) as a Rapid and Accurate Antimicrobial Susceptibility Testing Method for Staphylococcus aureus. EBioMedicine 4:95–103.

12. Lamsa A, Lopez-Garrido J, Quach D, Riley EP, Pogliano J, Pogliano K. 2016. Rapid Inhibition Profiling in Bacillus subtilis to Identify the Mechanism of Action of New Antimicrobials. ACS Chem Biol 11:2222–2231.

13. Htoo HH, Brumage L, Chaikeeratisak V, Tsunemoto H, Sugie J, Tribuddharat C, Pogliano J, Nonejuie P. 2019. Bacterial Cytological Profiling as a Tool To Study Mechanisms of Action of Antibiotics That Are Active against Acinetobacter baumannii. Antimicrob Agents Chemother 63:1–11.

14. Team RC. 2014. R: A language and environment for statistical computing. R Core Team (2014) R A Lang Environ Stat Comput R Found Stat Comput Vienna, Austria (available from http://www.R-project.org/).

15. Zhang J-H, Chung TDY, Oldenburg KR. 1999. A Simple Statistical Parameter for Use in Evaluation and Validation of High Throughput Screening Assays. J Biomol Screen 4:67–73.

16. Di Martino P, Cafferini N, Joly B, Darfeuille-Michaud A. 2003. Klebsiella pneumoniae type 3 pili facilitate adherence and biofilm formation on abiotic surfaces. Res Microbiol 154:9–16.

17. Hassan MZ, Sturm-Ramirez K, Rahman MZ, Hossain K, Aleem MA, Bhuiyan MU, Islam MM, Rahman M, Gurley ES. 2019. Contamination of hospital surfaces with respiratory pathogens in Bangladesh. PLoS One 14:e0224065.

18. Carpenter AE, Jones TR, Lamprecht MR, Clarke C, Kang IH, Friman O, Guertin DA, Chang JH, Lindquist RA, Moffat J, Golland P, Sabatini DM. 2006. CellProfiler: image analysis software for identifying and quantifying cell phenotypes. Genome Biol 7:R100.

19. Stringer C, Wang T, Michaelos M, Pachitariu M. 2020. Cellpose: a generalist algorithm for cellular segmentation. Nat Methods https://doi.org/10.1038/s41592-020-01018-x.

20. Van Laar TA, Chen T, You T, Leung KP. 2015. Sublethal concentrations of carbapenems alter cell morphology and genomic expression of Klebsiella pneumoniae biofilms. Antimicrob Agents Chemother 59:1707–1717.

21. Horii T, Kobayashi M, Sato K, Ichiyama S, Ohta M. 1998. An in-vitro study of carbapenem-induced morphological changes and endotoxin release in clinical isolates of gram-negative bacilli. J Antimicrob Chemother 41:435–442.

22. Zimmerman SB. 2006. Shape and compaction of Escherichia coli nucleoids. J Struct Biol 156:255–261.

23. Zahller J, Stewart PS. 2002. Transmission electron microscopic study of antibiotic action on Klebsiella pneumoniae biofilm. Antimicrob Agents Chemother 46:2679–2683.

24. Roth BL, Poot M, Yue ST, Millard PJ. 1997. Bacterial viability and antibiotic susceptibility testing with SYTOX green nucleic acid stain. Appl Environ Microbiol 63:2421–2431.

25. McKenzie K, Maclean M, Grant MH, Ramakrishnan P, MacGregor SJ, Anderson JG. 2016. The effects of 405 nm light on bacterial membrane integrity determined by salt and bile tolerance assays, leakage of UV-absorbing material and SYTOX green labelling. Microbiol (United Kingdom) 162:1680–1688.

26. Ang MLT, Pethe K. 2016. Contribution of high-content imaging technologies to the development of anti-infective drugs. Cytom Part A 89:755–760.

27. Zahir T, Camacho R, Vitale R, Ruckebusch C, Hofkens J, Fauvart M, Michiels J. 2019. High-throughput time-resolved morphology screening in bacteria reveals phenotypic responses to antibiotics. Commun Biol 2:269.

