## Supplementary data for "High-content imaging to phenotype antimicrobial effects on individual bacteria at scale"

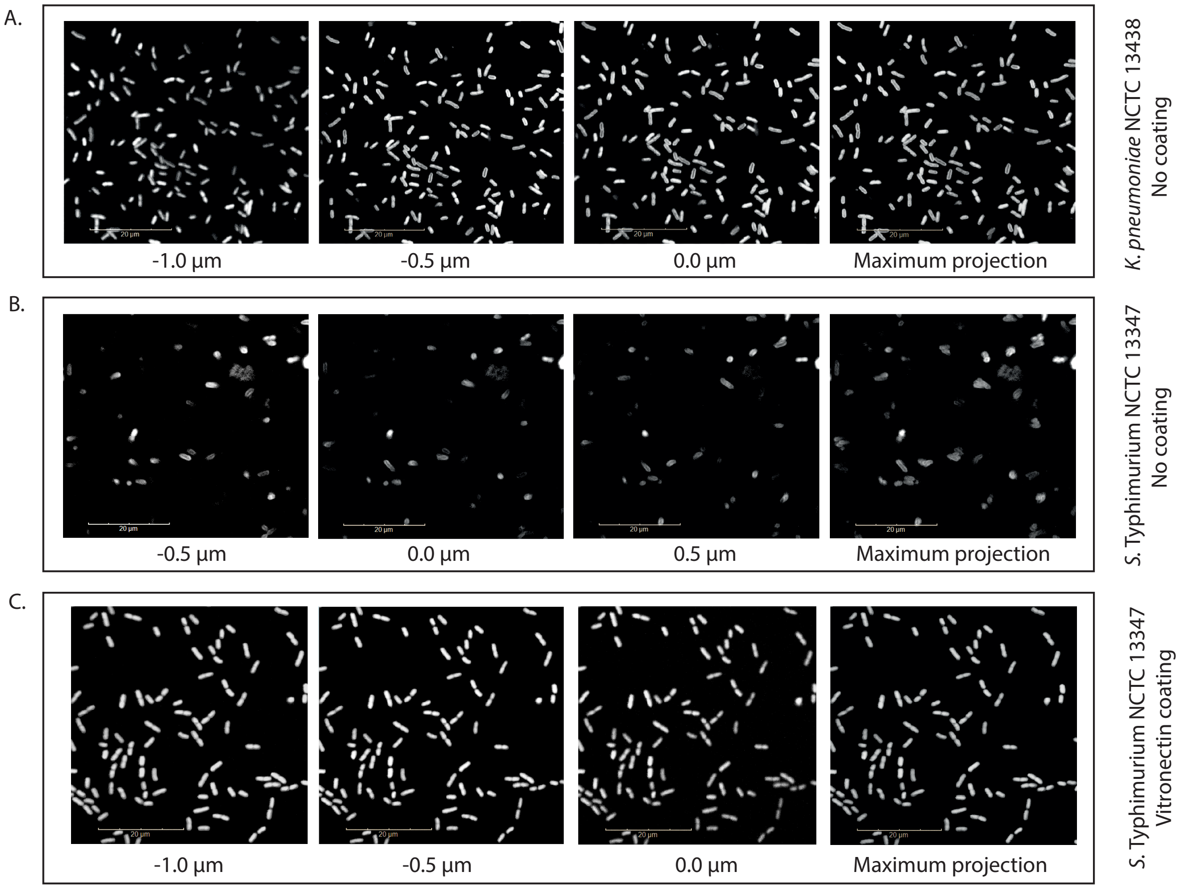

**Figure S1: Adhesion properties affect image quality.** Confocal images of *K. pneumoniae* isolate NCTC 13438 on a non-coated plate (A) and *S.* Typhimurium isolate NCTC 13347 on both non-coated (B) and vitronectin coated (C) plates. The first 3 images in each row represent different Z-stacks of the same field at 0.5 µm intervals, with the composite maximum projection image in the final column.

A.

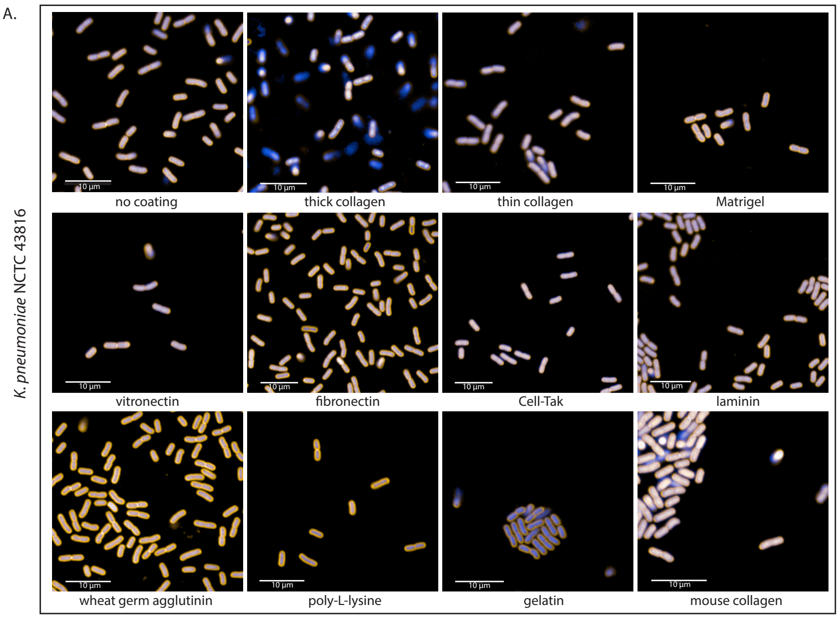
K. pneumoniae NCTC 43816
no coating
thick collagen
thin collagen
Matrigel
vitronectin
fibronectin
Cell-Tak
laminin
wheat germ agglutinin
poly-L-lysine
gelatin
mouse collagen

B.

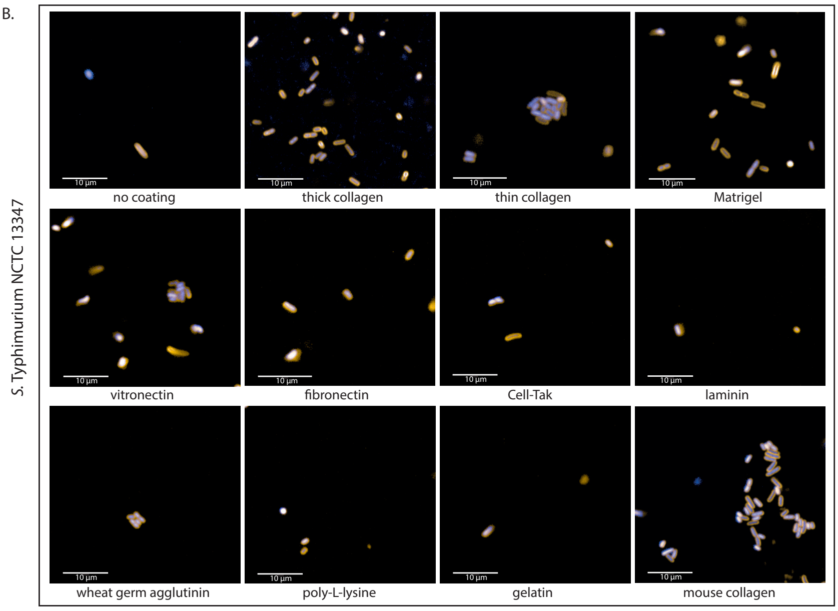
S. Typhimurium NCTC 13347
no coating
thick collagen
thin collagen
Matrigel
vitronectin
fibronectin
Cell-Tak
laminin
wheat germ agglutinin
poly-L-lysine
gelatin
mouse collagen

C.

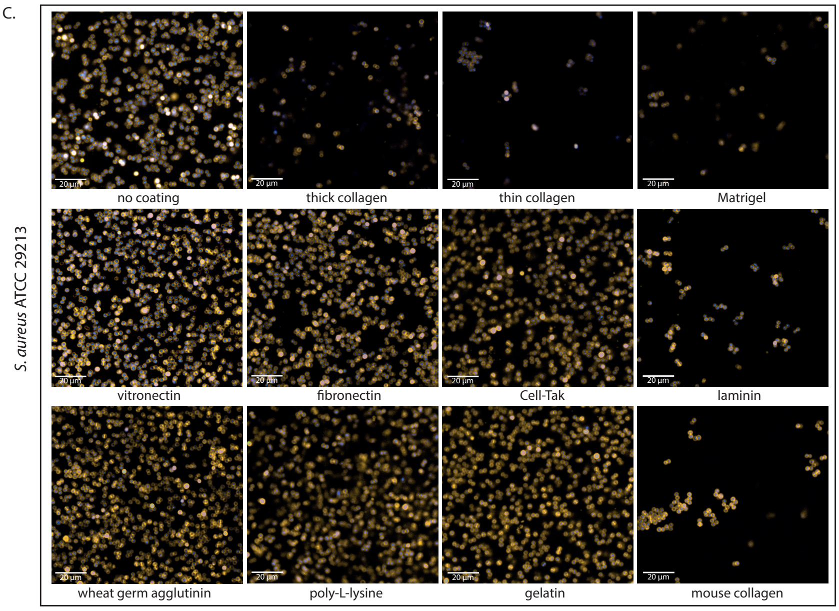
S. aureus ATCC 29213
no coating
thick collagen
thin collagen
Matrigel
vitronectin
fibronectin
Cell-Tak
laminin
wheat germ agglutinin
poly-L-lysine
gelatin
mouse collagen

**Figure S2: Representative images of isolates on different well coating matrices.** *K. pneumoniae* NCTC 43816 (**A**), *S.* Typhimurium NCTC 13347 (**B**), and *S. aureus* ATCC 29213 (**C**) were grown in non-coated wells or wells containing collagen, Matrigel, vitronectin, fibronectin, Cell-Tak, laminin wheat germ agglutinin poly-L-lysine, gelatin or mouse collagen for 2 hours, then fixed and stained with FM4-64 (membrane) and DAPI (nucleic acid). Images were acquired on the Opera Phenix using a 63x water immersion lens.

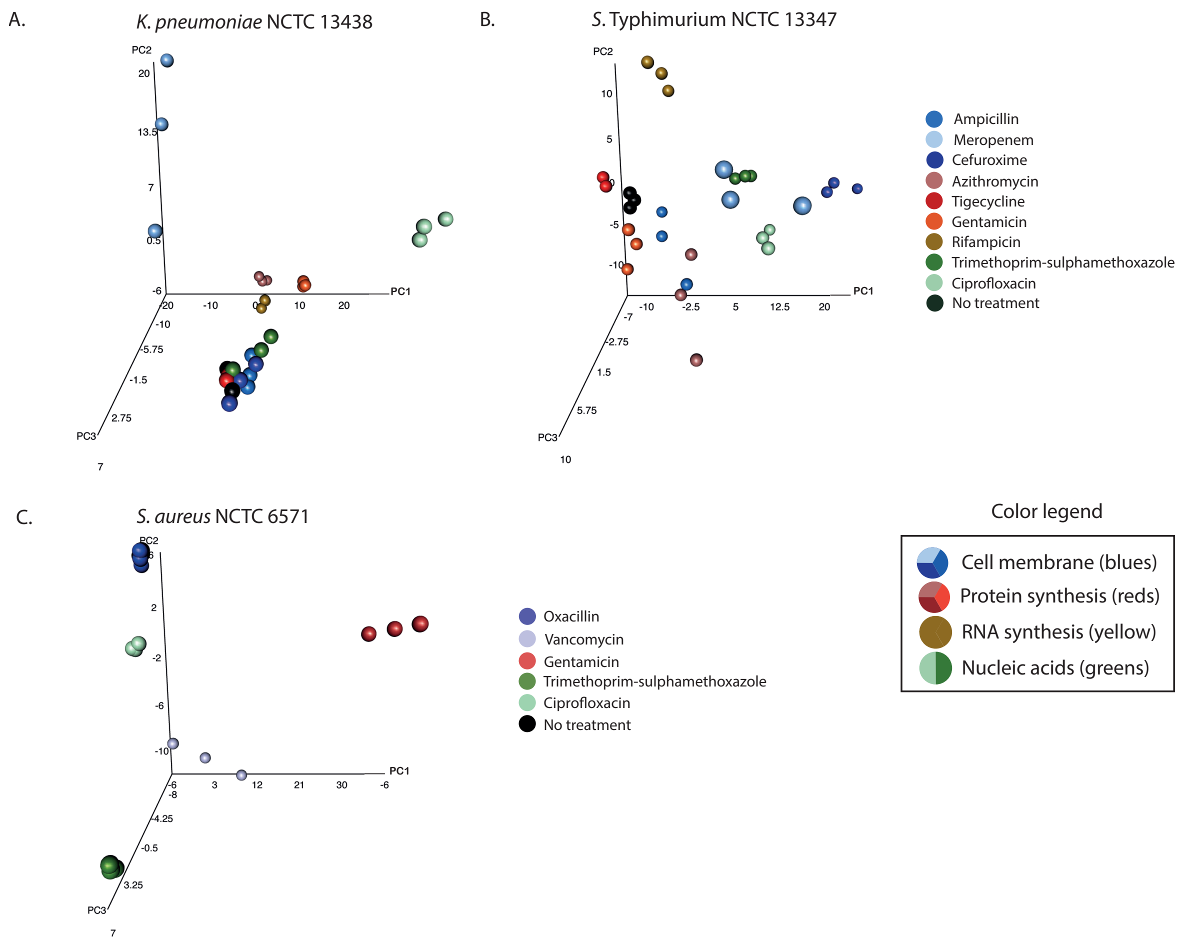

**Figure S3: PCA plots of bacteria under antimicrobial pressure.** Three-dimensional principal component analysis of the mean and standard deviation values of 62 morphological properties measured for single bacterial cells in each well for *K. pneumoniae* NCTC 13438 (**A**), *S.* Typhimurium NCTC 13347 (**B**), *S. aureus* NCTC 6571 (**C**) after 2hours exposure to antimicrobials. Technical triplicate repeats are shown.

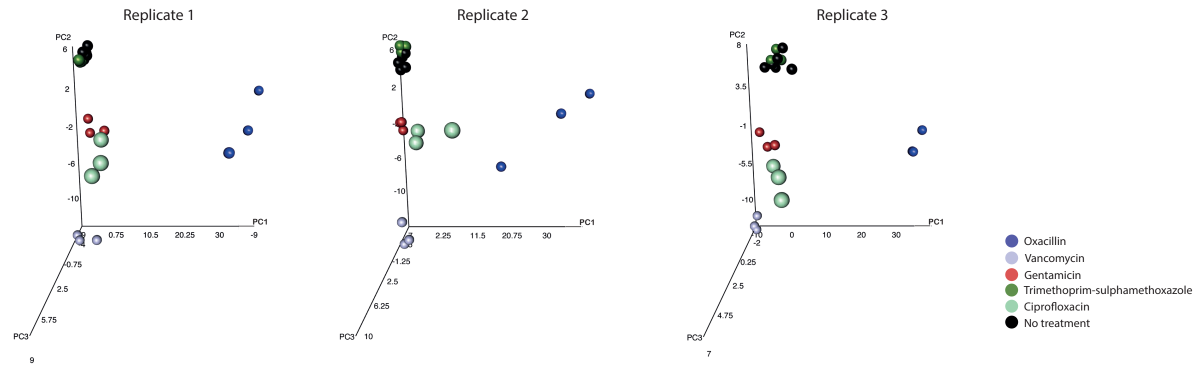

**Figure S4: Assay reproducibility.** Three-dimensional principal component analysis of the mean and standard deviation values of 62 morphological properties measured for single bacterial cells in each well for *S. aureus* ATCC 29213 after 2 hours exposure to antimicrobials. Three biological replicates with 3 technical replicates per assay are shown.

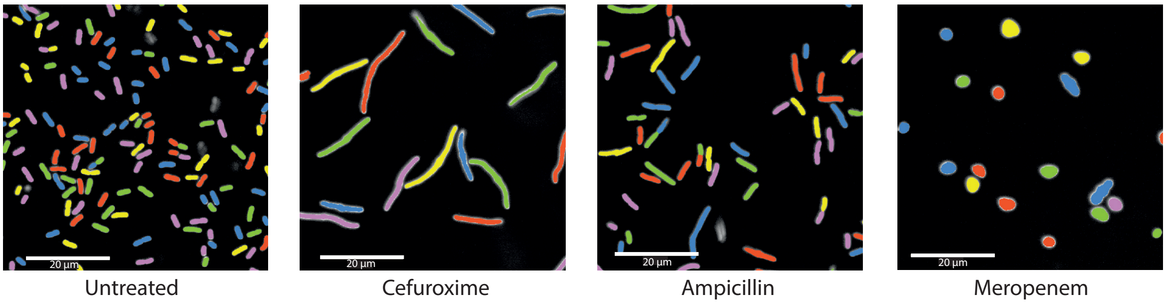

**Figure S5: Image segmentation is robust across different morphologies.** The Gram-negative analysis pipeline was applied to images of *K. pneumoniae* treated with cefuroxime, ampicillin and meropenem compared to an untreated control to illustrate the ability of the software to segment and identify individual bacteria with distinct morphologies. The different colours are used to highlight what has been selected as single bacteria.

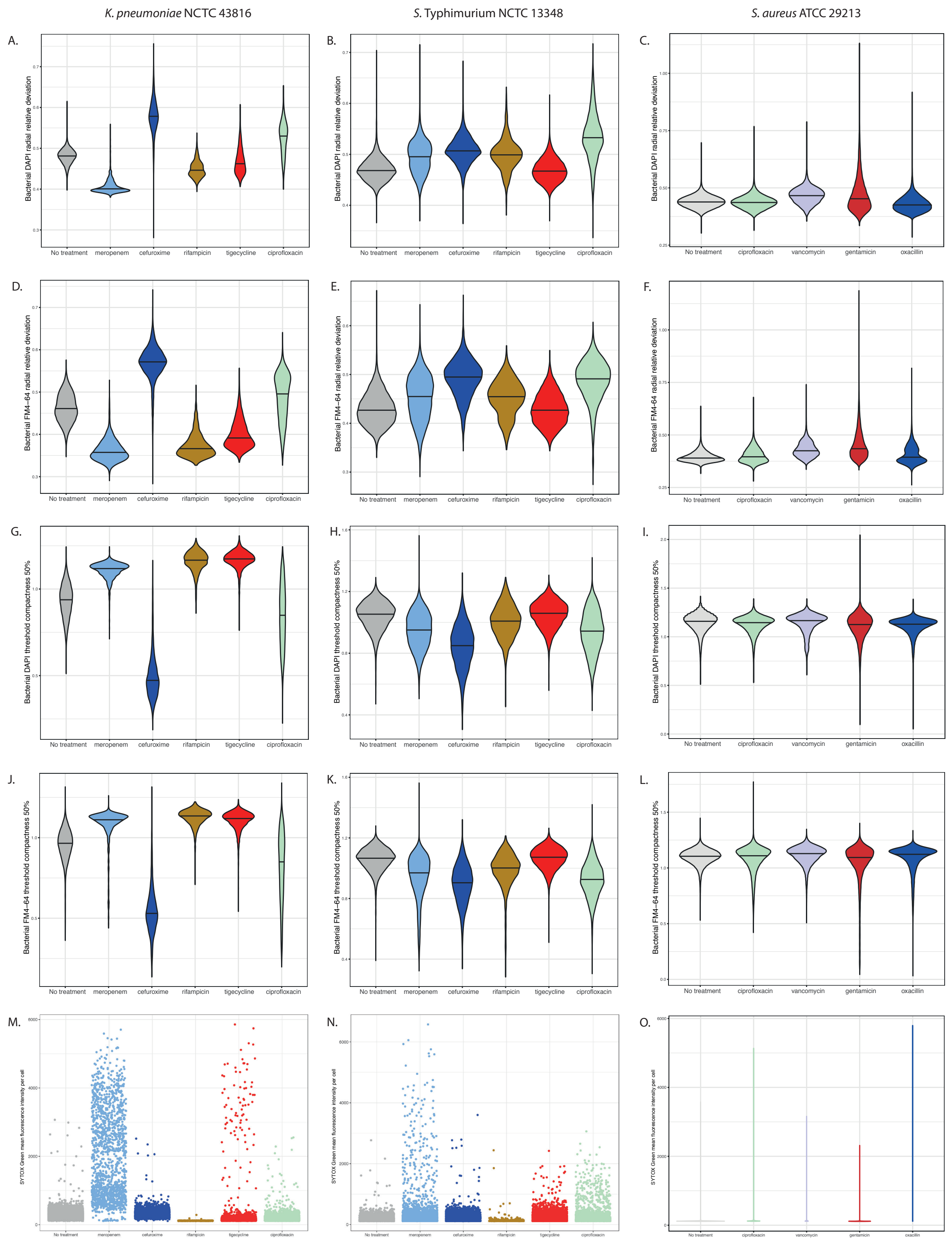

**Figure S6: Comparison of individual intensity-based morphological measurements.** Violin plots of bacterial DAPI radial relative deviation (A-C), bacterial FM4-64 radial relative deviation (D-F), bacterial DAPI threshold compactness 50% (G-I) and bacterial FM4-64 threshold compactness 50% (J-L) comparing *K. pneumoniae* NCTC 43816 and *S.* Typhimurium NCTC 13348 treated with meropenem, cefuroxime, rifampicin, tigecycline and ciprofloxacin, and *S. aureus* ATCC29213 treated with ciprofloxacin, vancomycin, gentamicin and oxacillin, with untreated controls. M-N shows scatter plots, and O a violin plot of SYTOX green intensity under the same conditions.

**Table S1: Plate coating conditions tested**

| Condition | Manufacturer | Catalogue Number | Final Concentration | Diluent | Number of washes | Wash buffer |
|---|---|---|---|---|---|---|
| Non-coated | N/A | N/A | N/A | N/A | N/A | N/A |
| Thick Collagen | Thermo Fisher | A10483 | 2mg/ml | $H_2O$, 1N NaOH, 10x PBS | 1 | PBS |
| Thin Collagen | Thermo Fisher | A10483 | 50µg/ml | 20mM Acetic Acid | 3 | PBS |
| Matrigel | Corning | 536231 | 0.5mg/ml | PBS | 1 | PBS |
| Vitronectin | Stem Cell Technologies | 7180 | 0.01mg/ml | Dilution buffer | 1 | Dilution buffer |
| Fibronectin | Thermo Fisher | 354008 | 50µg/ml | PBS | 1 | H2O |
| Cell-Tak | Corning | 354240 | 3.5mg/ml | 0.1N Sodium bicarbonate | 1 | H2O |
| Laminin | Sigma | L4544 | 5µg/ml | HBSS | 1 | HBSS |

### Supplementary Table S2: Gram-negative rods analysis pipeline (Harmony v4.9)

| | | | |
|---|---|---|---|
| **Input Image** | Input | | |
| | **Flatfield Correction**: Basic<br>Brightfield Correction<br>**Stack Processing**: Maximum Projection (*for non-adherent isolates, individual planes were analysed*)<br>**Min. Global Binning**: Dynamic | | |
| **Filter Image** | Input | Method | Output |
| | **Channel** : FM4-64 | **Method** : Texture SER<br>Filter : SER Ridge<br>Scale : 1 px<br>Normalization by : Kernel | Output Image : SER Ridge |
| **Find Image Region** | Input | Method | Output |
| | **Channel**: SER Ridge<br>**ROI**: None | Method : Common Threshold<br>Threshold : 0.4<br>Split into Objects<br>Area : > 100 px² | Output Population : Image Region<br>Output Region : Image Region |
| **Calculate Intensity Properties** | Input | Method | Output |
| | **Channel** : SER Ridge<br>**Population** : Image Region<br>**Region** : Image Region | **Method** : Standard<br>Mean | Property Prefix : Intensity Image Region SER Ridge |
| **Select Population** | Input | Method | Output |
| | **Population** : Image Region | **Method** Filter by Property<br>Intensity Image Region SER Ridge Mean : > 0.02 | Output Population : Image Region Selected |
| **Select Region** | Input | Method | Output |
| | **Population** : Image Region Selected<br>**Region** : Image Region | **Method** : Resize Region [µm/px]<br>Outer Border : -4µm<br>Restrictive Population : None<br>Restrictive Region :<br>Keep Image Border<br>Inner Border : INF µm | Output Population : Image Region Resized |
| **Select Region (2)** | Input | Method | Output |
| | **Population** : Image Region Selected<br>**Region** : Image Region Resized | **Method** : Standard<br>Border<br>Filled Region : INF µm² | Property Prefix : Image Region Resized Filled |
| **Select Region (3)** | Input | Method | Output |
| | **Population** : Image Region Selected<br>**Region** : Image Region Resized Filled | **Method** : Resize Region [µm/px]<br>Outer Border : 4 px<br>Restrictive Population : None<br>Restrictive Region :<br>Keep Image Border<br>Inner Border : INF px | Output Population : Image Region Resized Filled Resized |
| **Modify Population** | Input | Method | Output |

| | | | |
|---|---|---|---|
| | **Population :** Image Region Selected<br>Region : Image Region Resized Filled Resized | **Method** : Cluster by Distance<br>Distance : 0 px<br>Area : > 0 px² | Output Population : Modified Image Region Selected<br>Output Region : Modified Image Region |
| **Select Population (2)** | Input | Method | Output |
| | **Population** : Modified Image Region Selected | **Method** : Common Filters<br>Remove Border Objects<br>Region : Modified Image Region | Output Population : Modified Image Region Selected Border Removed |
| **Calculate Image** | Input | Method | Output |
| | | **Method** : By Formula<br>Formula : 100*(A-300)+100*(B-300)<br>Channel A : DAPI<br>Channel B : FM4-64<br>Negative Values : Set to Zero<br>Undefined Values : Set to Local Average | Output Image : Calculated Image |
| **Find Spots** | Input | Method | Output |
| | **Channel** : Calculated Image<br>**ROI** : Modified Image Region Selected Border Removed<br>**ROI Region** : Modified Image Region | **Method** : D<br>Detection Sensitivity : 0.5<br>Splitting Sensitivity : 0.1<br>Background Correction : 0.5<br>Calculate Spot Properties | Output Population : Spots |
| **Calculate Morphology Properties** | Input | Method | Output |
| | **Population** : Spots<br>**Region** : Spot | **Method** : Standard<br>Area<br>Roundness | Property Prefix : Spot |
| **Select Population (3)** | Input | Method | Output |
| | **Population** : Spot | **Method** Filter by Property<br>Spot Area [px²] : > 1 | Output Population : bacteria |
| **Calculate Morphology Properties (2)** | Input | Method | Output |
| | **Population** : bacteria<br>**Region** : Spot | **Method** : Standard<br>Area<br>Roundness<br>Width<br>Length<br>Ratio Width to Length | Property Prefix : bacteria |
| **Calculate Morphology Properties (3)** | Input | Method | Output |
| | **Population** : bacteria<br>**Region** : Spot | **Method** : STAR<br>Channel : DAPI<br>Symmetry<br>Threshold Compactness<br>Axial<br>Radial<br>Profile | Property Prefix : Bacteria DAPI |

| | | | |
|---|---|---|---|
| | | Profile Width : 3 px<br>**Sliding Parabola**<br>Curvature : 10<br>**Texture SER**<br>Scale : 1 px<br>Normalization by : Kernel | |
| **Calculate Morphology Properties (4)** | Input | Method | Output |
| | **Population** : bacteria<br>**Region** : Spot | **Method** : STAR<br>Channel : FM4-64<br>Symmetry<br>Threshold Compactness<br>Axial<br>Radial<br>Profile<br>Profile Width : 3 px<br>**Sliding Parabola**<br>Curvature : 10<br>**Texture SER**<br>Scale : 1 px<br>Normalization by : Kernel | Property Prefix : Bacteria FM4-64 |
| **Calculate Morphology Properties (5)** | Input | Method | Output |
| | **Population** : bacteria<br>**Region** : Spot | **Method** : STAR<br>Channel : SYTOX green<br>Symmetry<br>Threshold Compactness<br>Axial<br>Radial<br>Profile<br>Profile Width : 3 px<br>**Sliding Parabola**<br>Curvature : 10<br>**Texture SER**<br>Scale : 1 px<br>Normalization by : Kernel | Property Prefix : Bacteria SYTOX green |
| **Calculate Intensity Properties (3)** | Input | Method | Output |
| | **Channel** : DAPI<br>**Population** : bacteria<br>**Region** : Spot | **Method** : Standard<br>Mean<br>Standard Deviation | Property Prefix : Intensity Spot DAPI |
| **Calculate Intensity Properties (4)** | Input | Method | Output |
| | **Channel** : FM4-64<br>**Population** : bacteria<br>**Region** : Spot | **Method** : Standard<br>Mean<br>Standard Deviation | Property Prefix : Intensity Spot FM4-64 green |
| **Calculate Intensity Properties (5)** | Input | Method | Output |

| | | | |
|---|---|---|---|
| | **Channel** : SYTOX green<br>**Population** : bacteria<br>**Region** : Spot | **Method** : Standard<br>Mean<br>Standard Deviation | Property Prefix : Intensity Spot SYTOX green |
| **Select Population (4)** | Input | Method | Output |
| | **Population** : bacteria | **Method** Linear Classifier<br>Number of Classes : 3<br>Relative Spot Intensity<br>Corrected Spot Intensity<br>Uncorrected Spot Peak Intensity<br>Spot Contrast<br>Spot Background Intensity<br>Spot Area [px²]<br>Region Intensity<br>Spot to Region Intensity<br>Spot Area [µm²]<br>Spot Roundness<br>bacteria Area [µm²]<br>bacteria Roundness<br>bacteria Width [µm]<br>bacteria Length [µm]<br>bacteria Ratio Width to Length<br>Bacteria DAPI Symmetry 02<br>Bacteria DAPI Symmetry 03<br>Bacteria DAPI Symmetry 04<br>Bacteria DAPI Symmetry 05<br>Bacteria DAPI Symmetry 12<br>Bacteria DAPI Symmetry 13<br>Bacteria DAPI Symmetry 14<br>Bacteria DAPI Symmetry 15<br>Bacteria DAPI Threshold Compactness 30%<br>Bacteria DAPI Threshold Compactness 40%<br>Bacteria DAPI Threshold Compactness 50%<br>Bacteria DAPI Threshold Compactness 60%<br>Bacteria DAPI Axial Small Length<br>Bacteria DAPI Axial Length Ratio<br>Bacteria DAPI Radial Mean<br>Bacteria DAPI Radial Relative Deviation<br>Bacteria DAPI Profile 1/2<br>Bacteria DAPI Profile 2/2<br>Bacteria FM4-64 Symmetry 02<br>Bacteria FM4-64 Symmetry 03<br>Bacteria FM4-64 Symmetry 04<br>Bacteria FM4-64 Symmetry 05<br>Bacteria FM4-64 Symmetry 12<br>Bacteria FM4-64 Symmetry 13 | Output Population A : Single Cells<br>Output Population B : Dividing Cells<br>Output Population C : Other |

| | | |
|---|---|---|
| | | Bacteria FM4-64 Symmetry 14<br>Bacteria FM4-64 Symmetry 15<br>Bacteria FM4-64 Threshold Compactness 30%<br>Bacteria FM4-64 Threshold Compactness 40%<br>Bacteria FM4-64 Threshold Compactness 50%<br>Bacteria FM4-64 Threshold Compactness 60%<br>Bacteria FM4-64 Axial Small Length<br>Bacteria FM4-64 Axial Length Ratio<br>Bacteria FM4-64 Radial Mean<br>Bacteria FM4-64 Radial Relative Deviation<br>Bacteria FM4-64 Profile 1/2<br>Bacteria FM4-64 Profile 2/2<br>Bacteria Sytox Symmetry 02<br>Bacteria Sytox Symmetry 03<br>Bacteria Sytox Symmetry 04<br>Bacteria Sytox Symmetry 05<br>Bacteria Sytox Symmetry 12<br>Bacteria Sytox Symmetry 13<br>Bacteria Sytox Symmetry 14<br>Bacteria Sytox Symmetry 15<br>Bacteria Sytox Threshold Compactness 30%<br>Bacteria Sytox Threshold Compactness 40%<br>Bacteria Sytox Threshold Compactness 50%<br>Bacteria Sytox Threshold Compactness 60%<br>Bacteria Sytox Axial Small Length<br>Bacteria Sytox Axial Length Ratio<br>Bacteria Sytox Radial Mean<br>Bacteria Sytox Radial Relative Deviation<br>Bacteria Sytox Profile 1/2<br>Bacteria Sytox Profile 2/2<br>Bacteria Sytox Symmetry 02 SP-Filter<br>Bacteria Sytox Symmetry 03 SP-Filter<br>Bacteria Sytox Symmetry 04 SP-Filter<br>Bacteria Sytox Symmetry 05 SP-Filter<br>Bacteria Sytox Symmetry 12 SP-Filter<br>Bacteria Sytox Symmetry 13 SP-Filter<br>Bacteria Sytox Symmetry 14 SP-Filter<br>Bacteria Sytox Symmetry 15 SP-Filter<br>Bacteria Sytox Threshold Compactness 30% SP-Filter<br>Bacteria Sytox Threshold Compactness 40% SP-Filter<br>Bacteria Sytox Threshold Compactness 50% SP-Filter<br>Bacteria Sytox Threshold Compactness 60% SP-Filter<br>Bacteria Sytox Axial Small Length SP-Filter<br>Bacteria Sytox Axial Length Ratio SP-Filter<br>Bacteria Sytox Radial Mean SP-Filter<br>Bacteria Sytox Radial Relative Deviation SP-Filter |

| | | |
|---|---|---|
| | | Bacteria Sytox Radial Mean Ratio SP-Filter<br>Bacteria Sytox Profile 1/2 SP-Filter<br>Bacteria Sytox Profile 2/2 SP-Filter<br>Intensity Spot SYTOX green Mean<br>Intensity Spot SYTOX green StdDev |

**Method** : List of Outputs
**Population : bacteria**
Number of Objects
Intensity Spot DAPI Mean : Mean
Intensity Spot FM4-64 Mean : Mean
Intensity Spot SYTOX green Mean : Mean

**Population : Single Cells**
Number of Objects
Relative Spot Intensity : Mean+StdDev
Corrected Spot Intensity : Mean+StdDev
Uncorrected Spot Peak Intensity : Mean+StdDev
Spot Contrast : Mean+StdDev
Spot Background Intensity : Mean+StdDev
Spot Area [px²] : Mean+StdDev
Region Intensity : Mean+StdDev
Spot to Region Intensity : Mean+StdDev
Spot Area [µm²] : Mean+StdDev
Spot Roundness : Mean+StdDev
bacteria Area [µm²] : Mean+StdDev
bacteria Roundness : Mean+StdDev
bacteria Width [µm] : Mean+StdDev
bacteria Length [µm] : Mean+StdDev
bacteria Ratio Width to Length : Mean+StdDev
STAR Properties (Bacteria DAPI) : Mean+StdDev
STAR Properties (Bacteria FM4-64) : Mean+StdDev
STAR Properties (Bacteria Sytox) : Mean+StdDev
Intensity Spot DAPI Mean : Mean+StdDev
Intensity Spot FM4-64 Mean : Mean+StdDev
Intensity Spot SYTOX green Mean : Mean+StdDev
Intensity Spot SYTOX green StdDev : Mean+StdDev
Regression A-B : Mean+StdDev

**Population : Dividing Cells**
Number of Objects
**Population : Other**
Number of Objects

### Supplementary Table S3: Gram-positive cocci analysis pipeline (Harmony v4.9)

| | | | |
|---|---|---|---|
| **Input Image** | Input | | |
| | **Flatfield Correction**: Basic<br>Brightfield Correction<br>**Stack Processing**: Maximum Projection<br>**Min. Global Binning**: Dynamic | | |
| **Calculate Image** | Input | Method | Output |
| | | **Method** : By Formula<br>Formula : (100*(A-300))+(100*(B-300))<br>Channel A : FM4-64<br>Channel B : DAPI<br>Negative Values : Set to Zero<br>Undefined Values : Set to Local Average | Output Image : Calculated Image |
| **Find Image Region** | Input | Method | Output |
| | **Channel**: Calculated Image<br>**ROI**: None | Method : Common Threshold<br>Threshold : 0.56<br>Split into Objects<br>Area : > 0 px² | Output Population : Image Region<br>Output Region : Image Region |
| **Calculate Image (2)** | Input | Method | Output |
| | | **Method** : By Formula<br>Formula : 100*(A-300)<br>Channel A : DAPI<br>Negative Values : Set to Zero<br>Undefined Values : Set to Local Average | Output Image : Calculated Image (2) |
| **Find Spots** | Input | Method | Output |
| | **Channel** : DAPI<br>**ROI** : Image Region<br>**ROI Region** : Image Region | **Method** : D<br>Detection Sensitivity : 0.8<br>Splitting Sensitivity : 0.122<br>Background Correction : 0.5<br>Calculate Spot Properties | Output Population : Spots |
| **Select Region** | Input | Method | Output |
| | **Population** : Spots<br>**Region** : Spot | **Method** : Resize Region [µm/px]<br>Outer Border : -95 µm<br>Restrictive Population : Image Region<br>Restrictive Region : Image Region<br>Inner Border : INF µm | Output Population : Spots Resized |
| **Calculate Morphology Properties** | Input | Method | Output |
| | **Population** : Spots<br>**Region** : Spot Resized | **Method** : Standard<br>Area<br>Roundness | Property Prefix : Spot Resized |
| **Select Population** | Input | Method | Output |
| | **Population** : Spots | **Method** : Filter by Property<br>Spot Resized Area [µm²] : > 0.5 | Output Population : bacteria |

| | | | |
|---|---|---|---|
| **Calculate Morphology Properties (2)** | Input | Method | Output |
| | **Population :** bacteria<br>**Region** : Spot Resized | **Method** : Standard<br>Area<br>Roundness<br>Width<br>Length<br>Ratio Width to Length | Property Prefix : bacteria |
| **Calculate Morphology Properties (3)** | Input | Method | Output |
| | **Population** : bacteria<br>**Region** : Spot Resized | **Method** : STAR<br>Channel : DAPI<br>Symmetry<br>Threshold Compactness<br>Axial<br>Radial<br>Profile<br>Profile Width : 3 px<br>**Sliding Parabola**<br>Curvature : 10<br>**Texture SER**<br>Scale : 1 px<br>Normalization by : Kernel | Property Prefix : Bacteria DAPI |
| **Calculate Morphology Properties (4)** | Input | Method | Output |
| | **Population** : bacteria<br>**Region** : Spot Resized | **Method** : STAR<br>Channel : FM4-64<br>Symmetry<br>Threshold Compactness<br>Axial<br>Radial<br>Profile<br>Profile Width : 3 px<br>**Sliding Parabola**<br>Curvature : 10<br>**Texture SER**<br>Scale : 1 px<br>Normalization by : Kernel | Property Prefix : Bacteria FM4-64 |
| **Calculate Morphology Properties (5)** | Input | Method | Output |
| | **Population** : bacteria<br>**Region** : Spot Resized | **Method** : STAR<br>Channel : SYTOX green<br>Symmetry<br>Threshold Compactness<br>Axial | Property Prefix : Bacteria SYTOX green |

| | | | |
|---|---|---|---|
| | | Radial<br>Profile<br>Profile Width : 3 px<br>**Sliding Parabola**<br>Curvature : 10<br>**Texture SER**<br>Scale : 1 px<br>Normalization by : Kernel | |
| **Calculate Intensity Properties** | Input | Method | Output |
| | **Channel** : DAPI<br>**Population** : bacteria<br>**Region** : Spot Resized | **Method** : Standard<br>Mean | Property Prefix : Intensity Spot Resized DAPI |
| **Calculate Intensity Properties (2)** | Input | Method | Output |
| | **Channel** : FM4-64<br>**Population** : bacteria<br>**Region** : Spot Resized | **Method** : Standard<br>Mean | Property Prefix : Intensity Spot Resized FM4-64 |
| **Calculate Intensity Properties (3)** | Input | Method | Output |
| | **Channel** : SYTOX green<br>**Population** : bacteria<br>**Region** : Spot Resized | **Method** : Standard<br>Mean | Property Prefix : Intensity Spot Resized SYTOX green |
| **Select Population (2)** | Input | Method | Output |
| | **Population** : bacteria | **Method** : Common Filters<br>Remove Border Objects<br>Region : Spot | Output Population : bacteria Selected |

**Method** : List of Outputs
**Population** : bacteria
Number of Objects

**Population : bacteria Selected**
Number of Objects
Relative Spot Intensity : Mean+StdDev
Corrected Spot Intensity : Mean+StdDev
Uncorrected Spot Peak Intensity : Mean+StdDev
Spot Contrast : Mean+StdDev
Spot Background Intensity : Mean+StdDev
Spot Area [px²] : Mean+StdDev
Region Intensity : Mean+StdDev
Spot to Region Intensity : Mean+StdDev
Spot Resized Area [µm²] : Mean+StdDev
Spot Resized Roundness : Mean+StdDev
bacteria Area [µm²] : Mean+StdDev
bacteria Roundness : Mean+StdDev
bacteria Width [µm] : Mean+StdDev
bacteria Length [µm] : Mean+StdDev
bacteria Ratio Width to Length : Mean+StdDev
STAR Properties (Bacteria DAPI) : Mean+StdDev
STAR Properties (Bacteria FM4-64) : Mean+StdDev
STAR Properties (Bacteria Sytox) : Mean+StdDev
Intensity Spot Resized SYTOX green Mean : Mean+StdDev
Intensity Spot Resized SYTOX green StdDev : Mean+StdDev
Intensity Spot Resized DAPI Mean : Mean+StdDev
Intensity Spot Resized FM4-64 Mean : Mean+StdDev

**Method** : Standard Output
bacteria - Relative Spot Intensity : Mean
Output Name : bacteria - Relative Spot Intensity - Mean per Well

**Object Results**
Population : bacteria : ALL
Population : Spots : None
Population : Image Region : None
Population : bacteria Selected : Use Selected Well Results
